# Nuclear Pct1 couples phosphatidylcholine synthesis with membrane biogenesis

**DOI:** 10.64898/2026.04.21.719842

**Authors:** Pawel K. Lysyganicz, Zenon Toprakcioglu, Jennifer Guo, Antonio D. Barbosa, Benjamin J. Jenkins, Koini Lim, Albert Koulman, Tuomas P. J. Knowles, Marcus K. Dymond, David B. Savage, Symeon Siniossoglou

## Abstract

Phosphatidylcholine (PC) is the major eukaryotic phospholipid and its synthesis must be homeostatically controlled to prevent excess membrane and organelle growth. Here we investigate how PC synthesis by the Kennedy pathway is coordinated with membrane biogenesis. In budding yeast, the rate-limiting enzyme Pct1 is nuclear and reversibly associates with the inner nuclear membrane (INM) in response to lipid packing defects caused by low PC. We show that the enzymes acting after Pct1 to generate PC remain at the endoplasmic reticulum (ER) during pathway activation. Relocating the final PC synthesis step to different endomembrane sites does not alter the kinetics of Pct1 release from the INM, indicating that newly made PC equilibrates with the INM rapidly. In contrast, elevated phosphatidic acid locks Pct1 at the INM, prevents pathway inactivation and drives nuclear/ER membrane proliferation. These results support a model in which nuclear Pct1 senses lipid imbalance while ER-localized enzymes supply PC; disrupting this homeostasis leads to uncontrolled membrane biogenesis.

## Introduction

Biological membranes are an essential component of life. Membranes define the boundaries of cells; compartmentalize biochemical machineries within organelles; and provide a scaffold for complex reactions to occur. A major constituent of eukaryotic membranes is phosphatidylcholine (PC) (van Meer et al., 2008). PC synthesis must, therefore, match the rate of organelle and cell proliferation to support the need for new membranes. PC is also an essential component of lipoproteins and lipid droplets (LDs), pulmonary surfactant, and bile, and as a result, disruption of its homeostasis in humans is linked to several pathologies (Tavasoli et al., 2020). Therefore, cells have evolved tight regulatory mechanisms to maintain PC levels.

Both yeasts and mammals synthesize *de novo* PC via two pathways. The first route is the CDP-choline, or Kennedy, pathway consisting of three sequential reactions: firstly, soluble choline (Cho) is phosphorylated, then cytidylated and finally condensed with diacylglycerol (DG) to make PC (Fig. 1A) (Fagone and Jackowski, 2013). An alternative route, referred to as the methylation pathway, involves the triple methylation of the phosphatidylethanolamine (PE) headgroup to PC (Fig. 1A) (Ridgway and Vance, 1987). In humans, most cell types rely primarily on the Kennedy pathway for PC production, with the liver being the only tissue where the methylation pathway makes a substantial contribution to cellular PC (∼30%) (van der Veen et al., 2017). *Saccharomyces cerevisiae* (hereafter called yeast) can survive with either pathway active, but combined inactivation of both is lethal (Boumann et al., 2006; Kodaki and Yamashita, 1989). Notably, although these routes can substitute each other, they generate different molecular species of PC, suggesting functional diversification (Boumann et al., 2003; Boumann et al., 2004).

**Figure 1.**
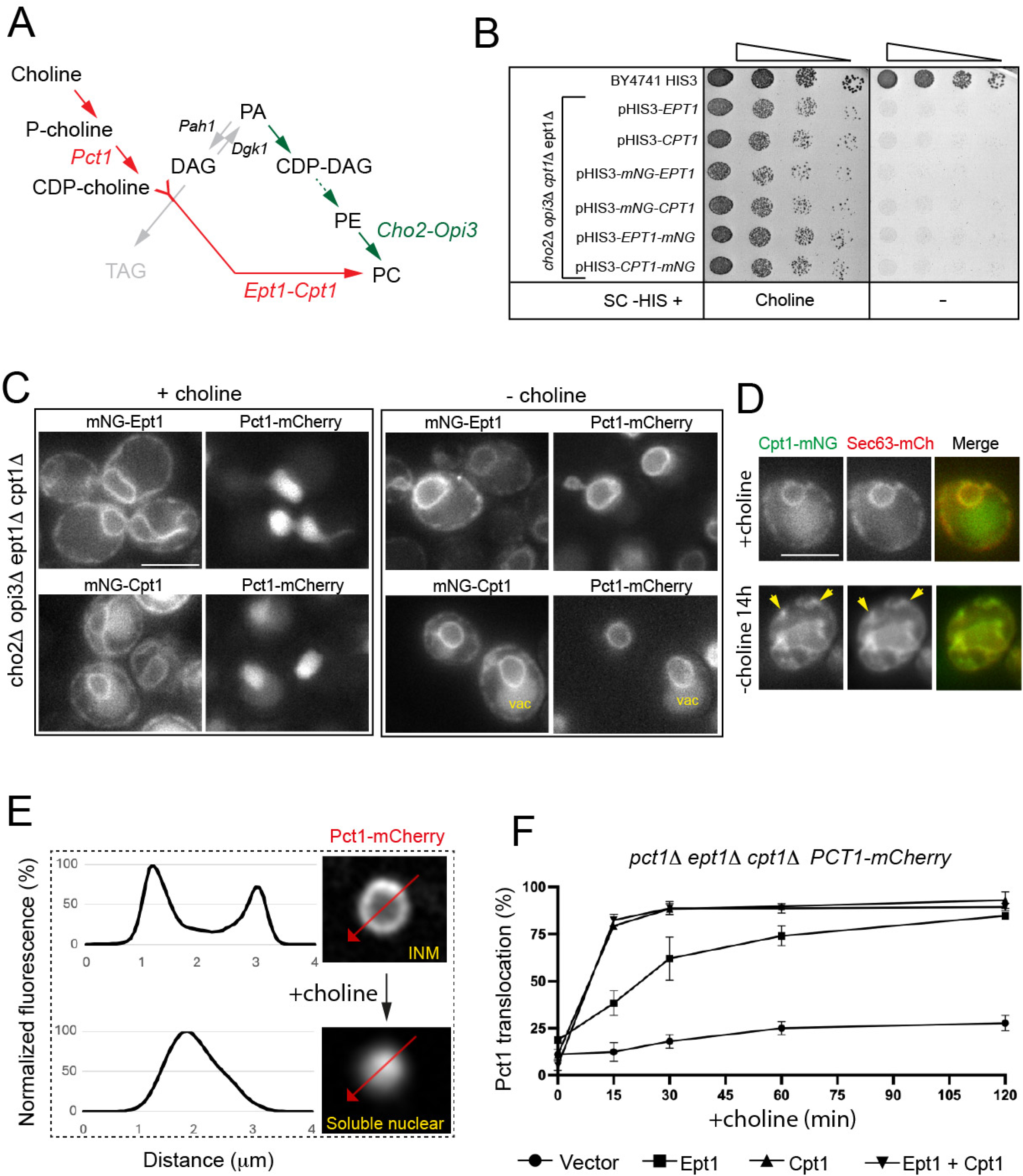
Nuclear translocation of Pct1 responds to the activity of ER-localized enzymes generating PC. (A) Graphical representation of the Kennedy (depicted in red) and methylation (in green) pathways; the key enzymes in each pathway are shown. (B) The indicated quadruple mutant (1PC) expressing the denoted plasmids, or a wild-type strain, were grown in synthetic media cultures, spotted on synthetic plates with or without 1mM choline and grown for two days at 30°C. (C) mNG-Cpt1 or mNG-Ept1 were expressed in 1PC expressing Pct1-mCh in the presence of 1mM choline, or following 8h choline depletion. (D) 1PC, expressing the denoted plasmids, were grown as in C with or without choline for 14h; arrowheads point to ER aggregates. (E) Representative pixel intensity profiles of wild-type cells expressing Pct1-mCh in the absence (“INM”) or presence (“soluble nuclear”) of choline; arrows indicate the direction of intensity quantification. (F) The *pct11 ept11 cpt11* mutant expressing Pct1-mCh and the denoted plasmid-borne genes, was grown in the absence of choline; following the addition of choline (1mM), the localization of Pct1-mCh was recorded at the indicated time-points and the rate of its translocation was calculated as shown in E (n=3).

The rate-limiting step of the Kennedy pathway is the formation of CDP-choline, catalysed by the CTP:phosphocholine cytidylyltransferase enzyme (CCTα; Pct1 in yeast) (Cornell and Ridgway, 2015). CCTα/Pct1 is highly conserved in eukaryotic kingdoms and binds reversibly to membranes via an amphipathic helix. Because PC is a cylindrical phospholipid that promotes formation of well-packed bilayers, insertion of the amphipathic helix is not favoured under PC-rich conditions. When PC levels are low, and membrane packing defects arise due to excess of conically shaped lipids like PE, or DAG, CCTα/Pct1 translocates onto membranes. This recruitment relieves an autoinhibitory interaction, inducing CCTα/Pct1’s catalytic activity and, consequently, PC synthesis (Cornell, 2020). These observations led to a model where low PC alters membrane physical properties; this is sensed by CCTα leading to changes in its activity in order to maintain PC homeostasis (Cornell, 2020; Cornell and Northwood, 2000). Hence, CCTα is referred to as PC “sensor”. Yeast Pct1 is confined to the nucleus (Haider et al., 2018; MacKinnon et al., 2009), where we have shown it partitions between an inactive nucleoplasmic pool and an active INM-bound state dictated by lipid packing defects (Haider et al., 2018). Nuclear localization of CCTα/Pct1 is conserved across diverse cell types (Cornell and Ridgway, 2015). Packing defects in LD monolayers can also recruit CCTα, including on nuclear LDs in hepatocytes (Soltysik et al., 2019) and cytoplasmic LDs in flies (Krahmer et al., 2011). In addition to packing defects, surface charge can also influence CCTα membrane recruitment (Cornell, 2016; Niu et al., 2024).

While the rate-limiting step of the Kennedy pathway is catalyzed by Pct1 at the INM, its CDP-choline product is used for the terminal reaction of PC synthesis that supports the growth of the endomembrane system in the cytoplasm. This spatial separation implies the need for inter-compartmental communication, through metabolite and lipid flux, to couple nuclear Pct1 activity to cytoplasmic PC production. Here, we sought to define how these processes are coordinated to ensure stable membrane growth, timely Pct1 inactivation once PC synthesis is induced, and the consequences of failing to reset this response.

## Results

### The enzymes catalyzing the terminal step of the Kennedy pathway localize at the ER membrane

The water-soluble product of Pct1, CDP-choline, is used by the integral membrane choline-phosphotransferase (Cpt1) to generate the final lipid product of the Kennedy pathway, PC. Ethanolamine-phosphotransferase (Ept1) can also utilize CDP-choline, as well as CDP-ethanolamine, to synthesize PC and PE, respectively (Hjelmstad and Bell, 1991; McMaster and Bell, 1994). To determine where this terminal step occurs, we first examined the localization of Cpt1 and Ept1. To ensure that our analysis was performed under conditions in which the Kennedy pathway is essential, we generated a quadruple deletion mutant lacking the terminal enzymes of both the methylation and Kennedy pathways (*cho21 opi31 ept11 cpt11,* hereafter called ΔPC) and confirmed that expression of either Cpt1 or Ept1 supports its growth in the presence of choline (Fig.1B and Fig. S1A). We then tagged Cpt1 and Ept1 with mNeonGreen either at their N-(mNG-Cpt1 and mNG-Ept1) or C-termini (Cpt1-mNG and Ept1-mNG), respectively and verified that all fusions were functional by growth complementation of ΔPC in the presence of choline (Fig.1B). In choline-replete conditions, all fusions displayed a characteristic ER pattern (Fig. 1C; Fig. S1B). Upon choline depletion, Pct1 relocalized from the nucleoplasm to the INM as expected, whereas Cpt1 and Ept1 showed no obvious redistribution and remained ER-associated (Fig. 1C). Prolonged choline depletion disrupted ER morphology and led to cortical ER membrane aggregates, where Cpt1-mNG co-localized (Fig. 1D). Finally, endogenously tagged Cpt1 and Ept1 in wild-type cells also localized to the ER membrane (Fig. S1C). Thus, the enzymes that catalyze the terminal step of PC synthesis in budding yeast remain ER-resident regardless of pathway activation state.

### Pct1 nuclear translocation tracks the activity of the terminal enzymes of the Kennedy pathway at the ER

Next, we sought to develop tools that would allow us to monitor PC synthesis in live cells using Pct1 as a reporter. We have previously shown that cellular PC deficiency results in the retention of Pct1 at the INM; and that supplementation of choline in media, which results in PC synthesis via the Kennedy pathway, causes the release of Pct1 – thereafter referred to as “translocation”-from the INM to the nucleoplasm (Haider et al., 2018). Line plot graphs of pixel intensities of Pct1-mCherry in these two locations are distinctly different (Fig. 1E). Based on these patterns, we developed an ImageJ-based workflow that assigns the location of Pct1-mCherry to the INM or the nucleoplasm (see Materials and Methods; Fig. 1E). To validate this approach, we applied it to quantify the translocation of Pct1-mCherry in wild-type or mutant cells compromised in PC synthesis via the Kennedy pathway. As expected, Pct1-mCherry failed to translocate to the nucleoplasm in cells lacking Ept1 and Cpt1 following the addition of choline; co-expression of Cpt1 and Ept1 together, or of Cpt1 alone, restored the maximal translocation of Pct1-mCherry, while expression of Ept1 alone resulted in a significantly slower rate of Pct1-mCherry translocation (Fig. 1F). These data are consistent with previous observations that Cpt1 is responsible for the majority of PC made via the CDP-choline pathway, while Ept1 makes a minor contribution (McMaster and Bell, 1994). Together, these results validate our ImageJ pipeline as a quantitative readout of Pct1 localization kinetics, providing a proxy for Kennedy-pathway activity *in vivo*.

### Kennedy pathway-derived PC redistributes rapidly across the endomembrane network

To investigate the significance of the spatial organization of the Kennedy pathway, we altered the terminal site of PC synthesis. We used the anchor-away technique (Haruki et al., 2008), which allows the conditional re-location of a protein of interest to a new subcellular compartment via rapamycin-dependent complex formation between FKBP12 and the FRB domain (Fig. 2A). We co-expressed Cpt1-FRB-GFP with FKBP12 fusions of either Pil1, a protein tightly associated with distinct domains of the plasma membrane, or Heh1, a component of the INM (Fig. 2A). In the absence of rapamycin, Cpt1-FRB-GFP displayed its typical ER distribution (Fig. 2B, top panels). Rapamycin addition drove Cpt1-FRB-GFP to bright cortical puncta (Pil1 anchor) or perinuclear rings (Heh1 anchor) with a concomitant loss of its peripheral ER localization (Fig. 2B).

**Figure 2.**
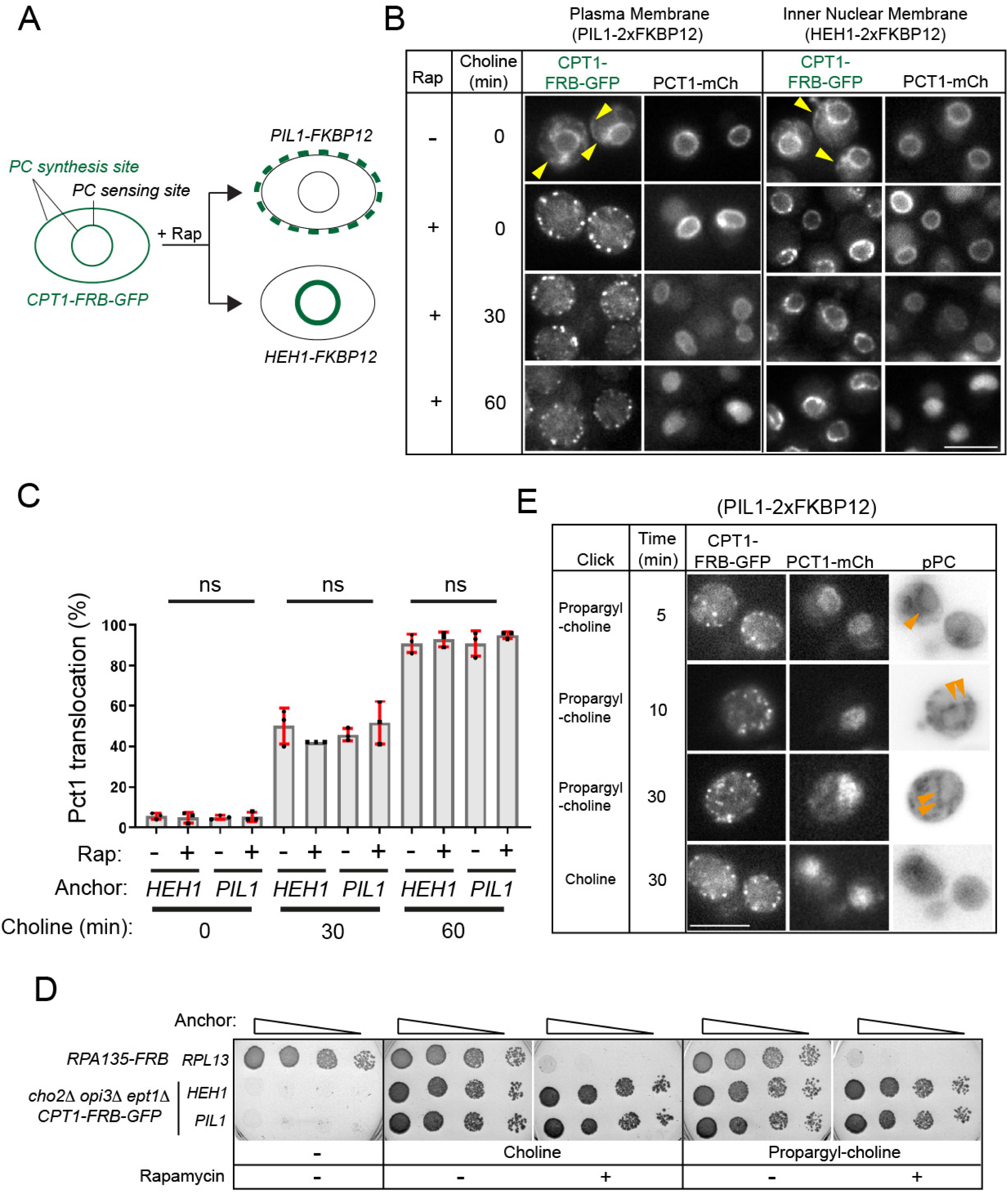
Pct1 nuclear translocation is independent of the site of PC synthesis. (A) Schematic of the anchor away method, which allows the relocation of Cpt1-FRB-GFP either to plasma membrane domains (PIL1-FKBP12) or the INM (HEH1-FKBP12). (B) A triple deletion mutant (*cho2Δ opi3Δ ept1Δ*) expressing the genomically integrated PIL1-or HEH1-FKBP12 anchors and Cpt1-FRB-GFP, and the plasmid-borne Pct1-mCh, were starved from choline for 7 hours; choline was then added for the indicated times, in the absence or presence of rapamycin (Rap); arrowheads indicate the cortical ER pool of Cpt1-FRB-GFP in the absence of rapamycin.(C) Quantification of Pct1 translocation shown in B; the anchor, the presence of rapamycin and the duration of choline supplementation, are denoted; data are means from three experiments +/- SD and analysed using a one-way ANOVA with Sidak’s post hoc test; ns, p≥ 0.05. (D) *cho2Δ opi3Δ ept1Δ* expressing the PIL1- or HEH1-FKBP12 anchors and Cpt1-FRB-GFP, were spotted on plates with the indicated supplements; cells expressing the essential RNA PolI subunit RPA135 as an FRB fusion were used as a control; cells were grown for three days at 30°C. (E) *ept1Δ* cells expressing the fusions shown, were treated with rapamycin (1h) to anchor Cpt1-FRB-GFP to the plasma membrane, followed by the addition of propargyl-choline, or choline, for the indicated times, fixed, and phosphatidyl-propargyl-choline (PC*) was then visualized by click-chemistry. PC* micrographs are shown as inverted grey-scale images; arrows indicate nuclear membranes containing PC*.

To monitor the impact of Cpt1 relocation on nuclear PC sensing, we followed Pct1 dynamics following choline addition. To ensure that Cpt1 is the sole source of PC synthesis, we performed these experiments in a strain lacking Cho2, Opi3 and Ept1. Remarkably, the movement of Pct1 remained unaffected regardless of whether Cpt1 is at the plasma membrane or at its native ER (Fig. 2C, compare +/- rapamycin within each anchor and time point). Moreover, contrary to our expectation that INM-anchored Cpt1 would promote faster translocation of Pct1, since PC would be produced locally at its sensing site, Pct1 translocation was similar in cells with Cpt1 anchored at the INM or the plasma membrane (Fig. 2C, compare the two anchors at each time-point). Consistently, Cpt1 anchored to the plasma membrane or INM, as the sole source of PC, rescued the growth of the *cho21 opi31 ept11* mutant, as effectively as when localizing to its native ER location (Fig. 2D). Similar anchoring experiments in a methylation-competent *CHO2 OPI3* background showed an overall faster Pct1 response to choline, as expected, but anchoring Cpt1 again did not alter the kinetics of Pct1 translocation (Fig. S2). Together, these observations suggest that the location of PC synthesis does not appreciably influence the rate of Pct1 inactivation in the nucleus.

These results suggest that PC made at the plasma membrane rapidly equilibrates throughout the cell and reaches the INM. To visualize PC directly, we applied a click-chemistry method to metabolically incorporate an alkyne-containing choline analogue, propargyl-choline (Pr-Cho), into phosphatidyl-propargyl-choline (pPC) (Iyoshi et al., 2014; Jao et al., 2009). Pr-Cho can functionally substitute Cho in the Kennedy pathway, because it sustains the growth of the *cho21 opi31 ept11* strain in the absence of choline (Fig. 2D). We then anchored Cpt1 to the plasma membrane and monitored the production of pPC by the addition of Pr-Cho. Even after a 5 min incubation with Pr-Cho, which is the earliest time point we could process the cells for the click reaction, a nuclear membrane labelling could be discerned (Fig. 2E), consistent with the observed dissociation of Pct1 in the nucleus. No labelling was detected in cells incubated with choline instead of Pr-Cho (Fig. 2E), or when Pr-Cho was added to cells lacking a functional Kennedy pathway (Fig. S3), confirming the specificity of the labelling. Collectively, these data indicate that the response of Pct1 is insensitive to whether PC is produced at the plasma membrane, INM or ER, and support a model in which once synthesized, PC is rapidly distributed within the cell.

### Kennedy pathway activation deforms the nucleus

Next, we asked whether the equilibration of newly synthesized PC across the ER–nuclear envelope network affects the organization of the nuclear membrane. Following choline addition, nuclei became deformed, as evidenced by the decrease in nuclear circularity (3A and 3B). This effect required a functional Kennedy pathway, as nuclear shape remained unchanged in *ept1*1 *cpt1*1 cells (Fig. 3A and 3B), indicating that is mediated by the production of new PC rather than the addition of choline. To follow the very early events after induction of PC synthesis, we monitored cells using a microfluidics device during choline addition (Fig. 3C and Materials and methods). Under these conditions, Pct1 translocated from the INM to the nucleoplasm within the first 10 minutes in 80% of cells (Fig. 3D and 3E). Consistent with our results in bulk cultures, nuclear circularity of cells immobilized within the microfluidics device also decreased following choline addition (Fig. 3F), and this nuclear deformation correlated with the release of Pct1 from the INM (Fig. 3G). Together, these findings are consistent with a model in which activation of the Kennedy pathway rapidly remodels nuclear shape.

**Figure 3.**
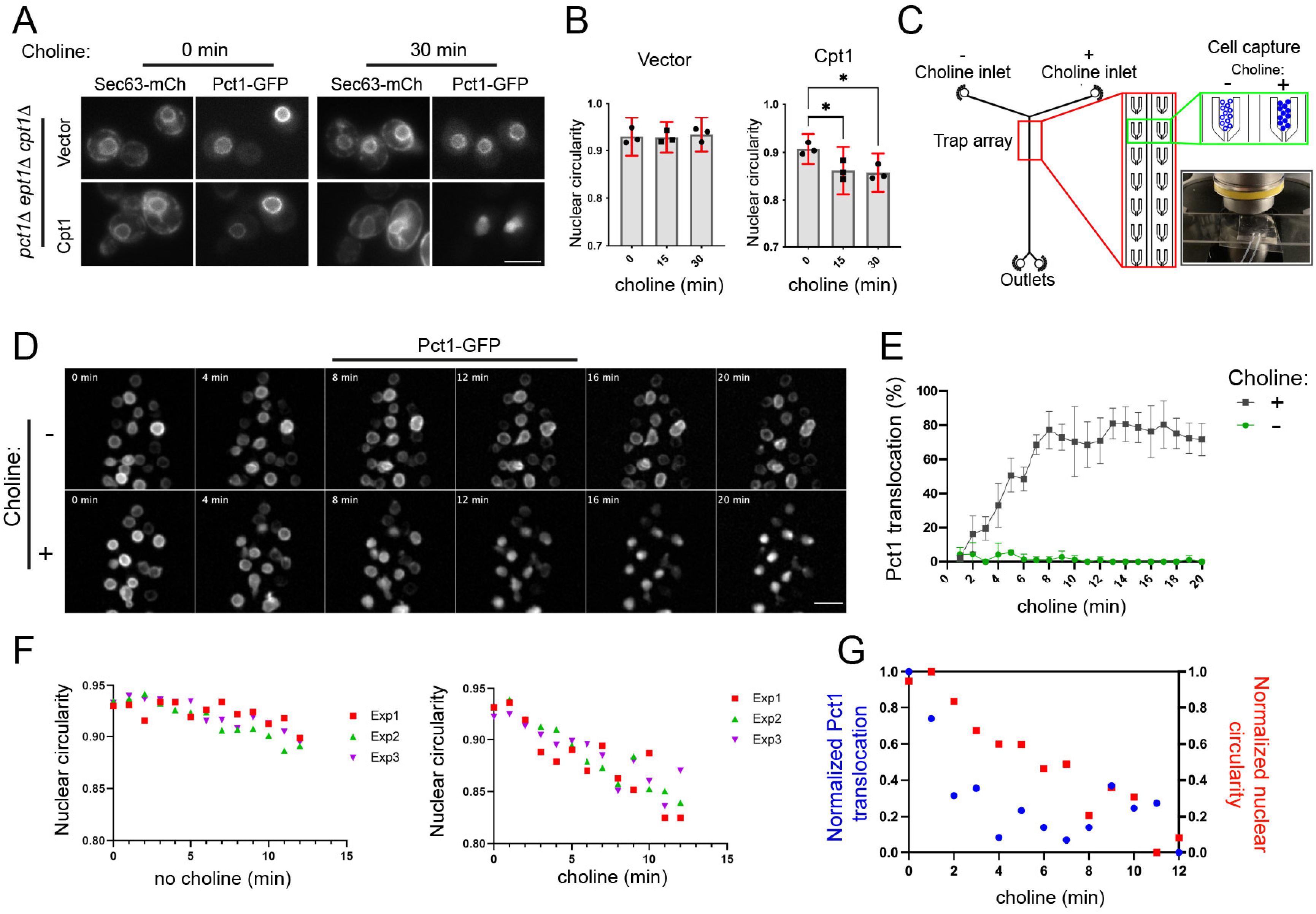
Choline supplementation deforms the nucleus in a Cpt1-dependent manner. (A) *pct11 ept11 cpt11* cells expressing the denoted reporters, and a vector or Cpt1, were treated with 1mM choline as indicated. (B) Quantification of changes in nuclear circularity in cells from (A) from three experiments; data are means from three experiments +/- SD and analysed using a one-way ANOVA with Sidak’s post hoc test; *p< 0.05. (C) Microfluidic device and cell trap setup (D) Representative time frames from a time-lapse movie recording the response of Pct1-mCh in microfluidic chamber-trapped cells, following the addition (lower panel), or not (upper panel) of 1mM choline. (E) Quantification of Pct1-mCh translocation from cells in microfluidic chamber-trapped cells, +/- choline (n=4 time-lapse movies). (F) Quantification of Pct1-mCh translocation from cells in microfluidic chamber-trapped cells; cells analyzed were co-expressing Sur4-GFP which was used to calculate nuclear circularity; data from three independent time-lapse movies are shown; left, no choline; right, 1mM choline. (G) Correlation of changes in nuclear shape and Pct1 movement; data are averages of circularity and Pct1 translocation taken from E, and have been normalized from 0 to 1.

### The activity of the PA phosphatase Pah1 is required for the release of Pct1 from the INM

PA is the central precursor of most phospholipid classes; DG, which can be derived from PA, is used with CDP-Cho generated by Pct1, to make PC (Fig. 1A). We therefore asked how the activity of the yeast lipin Pah1, which dephosphorylates PA to generate DG, influences Pct1 translocation and activity. Loss of Pah1 activity results in multiple defects, including the accumulation of PA, the upregulation of phospholipid synthesis and the expansion of the nuclear/ER membrane (Han et al., 2006; Santos-Rosa et al., 2005). We first confirmed that Pct1 still targets the INM in *pah1*1 cells by co-localizing it with Heh2, a member of the LEM (Lap2-Emerin-Man1) family and INM resident (King et al., 2006) (Fig. 4A). Following the addition of choline to *pah1*1 cells, we noticed that Pct1 remained membrane-bound even after 2 h (Fig. 4B). Supplementation of Pr-Cho to *pah1*1 led to ER membrane labelling, indicating that despite blocking PA to DG conversion in this mutant, the Kennedy pathway remains operational (Fig. 4C). We further tested how Pct1 responds in cells lacking Nem1, a phosphatase required for the activation of Pah1 (Santos-Rosa et al., 2005) (Fig. 4D); *nem1*1 cells display similar, albeit less severe, phenotypes when compared to *pah1*1 (Santos-Rosa et al., 2005). Pct1 remained bound to the INM for longer in *nem1*1 compared to wild-type cells after the addition of choline (Fig.4D). Notably, this change coincided with an exacerbation of the nuclear/ER membrane expansion of *nem1*1 cells after 2 h in the presence of choline (Fig. 4D). Taken together, these results indicate that deficiency in Pah1 activity disrupts the homeostatic regulation of Pct1 in response to choline supplementation; we therefore set out to determine the factors underlying this deregulation.

**Figure 4.**
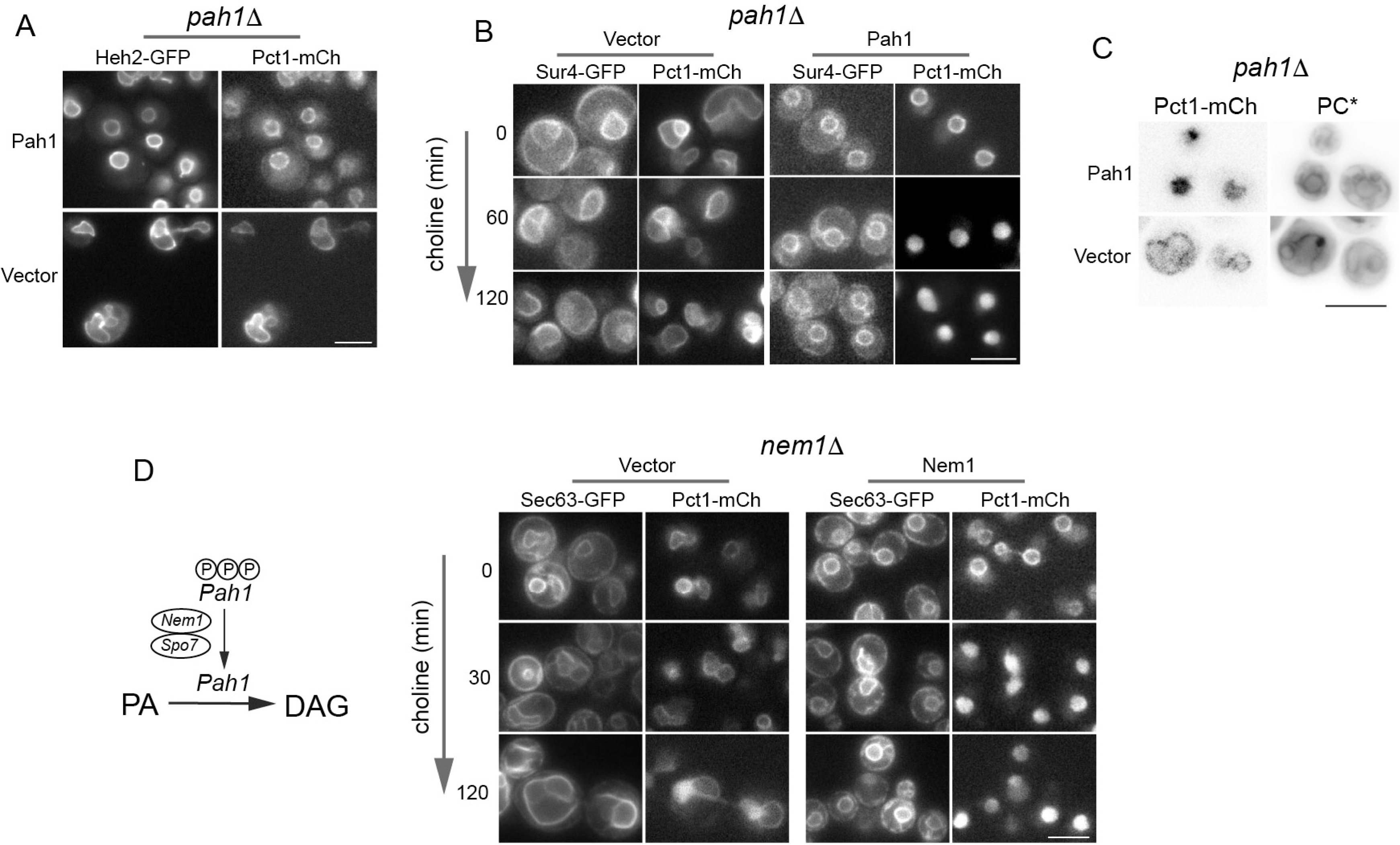
The activity of the PA phosphatase Pah1 is required for the release of Pct1 from the INM. (A) *pah1*1 cells, co-expressing Heh2-GFP and Pct1-mCh, and the indicated plasmids, were imaged. (B) *pah1*1 cells, co-expressing Pct1-mCh and the ER reporter Sur4-GFP, and the indicated plasmids, were supplemented with choline for the denoted times and imaged. (C) *pah1*1 cells, expressing Pct1-mCh and the indicated plasmids, were supplemented with propargyl-choline for 30 min, fixed, and phosphatidyl-propargyl-choline (PC*) was then visualized by click-chemistry. (D) Left panel: schematic of the role of Nem1-Spo7 in the activation of Pah1; right panel: *nem1*1 cells, expressing Sec63-GFP and Pct1-mCh, and the indicated plasmids were supplemented with choline for the denoted times and imaged. Scale bars in all micrographs, 5 μm.

### Constitutive membrane recruitment of Pct1 activity induces nuclear and ER expansion

The severe membrane defects in *pah1*1 make it difficult to discern the effects of Pct1 recruitment in these cells. We therefore sought to use a system that allows us to determine the consequences of Pct1 and Kennedy pathway activity in this mutant. We previously showed that many *pah1*1 defects are alleviated by deletion of the DG kinase Dgk1 (Han et al., 2008a; Han et al., 2008b) (Fig. 1A). Specifically, in *pah1*1 *dgk1*1 cells, elevated PA levels, derepressed phospholipid synthesis and nuclear/ER membrane proliferation are restored to near wild-type, levels. Therefore, we investigated the behaviour of Pct1 in the *pah1*1 *dgk1*1 mutant. As expected, similar to wild-type cells (Fig. 5A), *pah1*1 *dgk1*1 cells displayed normal ER morphology (Fig. 5B, 0 min); following the supplementation of choline, the nucleus and ER became initially deformed (30 min) and then underwent significant expansion, filling the cytoplasm with large membrane sheets (60 and 120 min, Fig. 5B). This membrane expansion correlated with the failure of Pct1 to dissociate from the INM in choline-replete media, in contrast to wild-type cells where Pct1 became intranuclear within 30 min (compare Pct1-GFP in Fig. 5A and B). Importantly, membrane expansion was blocked when Pct1 was removed (Fig. S4A), or when its ability to associate with the INM was disrupted by mutating four key hydrophobic residues within its amphipathic helix (Pct1-4M; Fig.5C). Pct1 and Pct1-4M were expressed at similar levels (Fig. 5D). Therefore, stable membrane binding of Pct1 and a functional Kennedy pathway are required for ER membrane expansion under these conditions.

**Figure 5.**
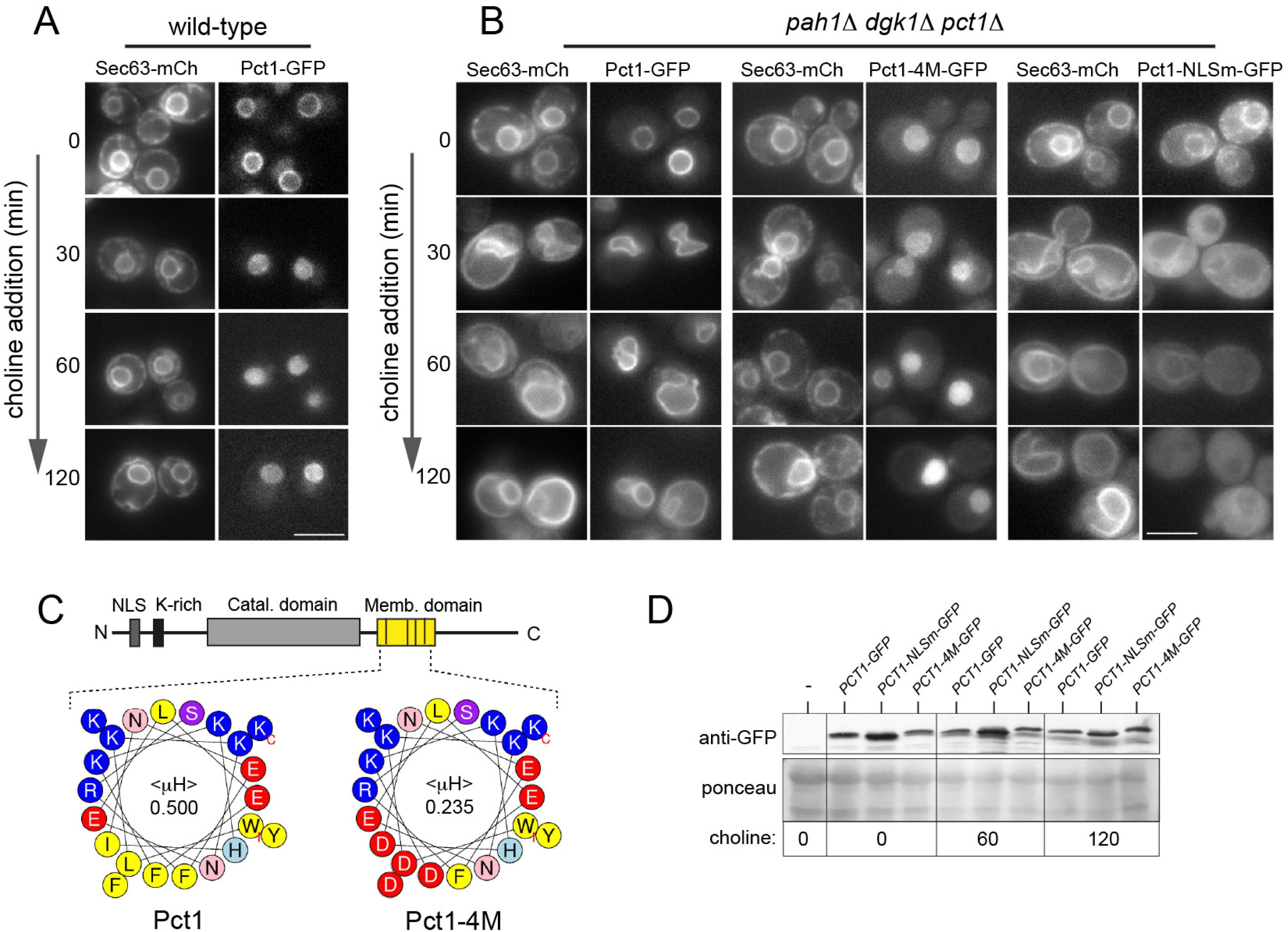
Constitutive binding of Pct1 at the INM drives membrane proliferation and nuclear/ER membrane expansion. (A) Wild-type cells or (B) *pah1*1 *dgk1*1 *pct1*1 cells, co-expressing Sec63-GFP and the denoted Pct1-mCh plasmids, were supplemented with choline for the indicated times, and imaged; scale bars, 5 μm. (C) Schematic of the primary structure of Pct1 with the position of the NLS, lysine-rich stretch and the membrane-binding domain indicated; the helical wheel plots of the amphipathic helix of the wild-type Pct1 and the Pct1-4M mutant and the hydrophobic moment values (μH), generated by Heliquest (Gautier et al., 2008), are shown; residues shown are 261 to 282; non-polar residues are shown in yellow. (D) Western blot of protein extracts from *pah1*1 *dgk1*1 *pct1*1 cells expressing the denoted Pct1 plasmids; lower panels show part of the ponceau dye-stained membrane.

We next asked whether Pct1 can drive membrane biogenesis from the cytoplasmic ER. Pct1 contains two short basic stretches at its N-terminus implicated in its nuclear import (Haider et al., 2018; MacKinnon et al., 2009); consistently, Pct1-NLSm carrying lysine to alanine mutations in these stretches, is found at the ER (Fig. 5B). Choline addition in Pct1-NLSm-expressing cells triggered membrane expansion indistinguishable from that seen with the wild-type Pct1 (Fig. 5B). Although Pct1-NLSm showed a progressive reduction in ER enrichment (Fig. 5B), membrane expansion was still abolished when the hydrophobic face of its amphipathic helix was mutated (Pct1-NLSm-4M; Fig. S4B), indicating that membrane binding remains essential for activity. Thus, Pct1 can promote membrane proliferation when recruited either to the INM or to cytoplasmic ER membranes, and failure to release Pct1 from membranes in choline-replete conditions locks the Kennedy pathway in an active state and drives membrane growth.

Because PA has been proposed to regulate CCTα and contribute to lipin-associated membrane expansion in other systems (Craddock et al., 2015; Yang et al., 2020; Zhang et al., 2019), we asked whether Kennedy pathway activity similarly underlies the membrane proliferation observed in *pah1*1 cells. Although Pah1/lipin inactivation is known to trigger membrane proliferation across diverse cell types and choline can drive the accumulation of nuclear/ER membrane sheets in our system, this mechanism cannot account for the canonical *pah1*1 phenotype in yeast: *pah1*1 *pct1*1 or *pah1*1 *ept1*1 *cpt1*1 cells, lacking a functional Kennedy pathway, display still membrane proliferation similar to that seen in *pah1*1 cells (Fig. S5). Thus, while Kennedy pathway hyperactivation can drive membrane biogenesis under conditions of elevated PA, the membrane expansion associated with loss of Pah1 in yeast must also involve Kennedy-independent mechanisms.

### Constitutive recruitment of Pct1 takes place at PA-enriched INM and induces phospholipid synthesis

Given that PA accumulation can both promote membrane proliferation and stabilize CCTα/Pct1 membrane binding, we asked whether PA increases at the INM under conditions where Pct1 remains membrane-bound. Therefore, we used a biosensor, consisting of a PA binding domain fused to a nuclear localization signal (Romanauska and Kohler, 2018), to determine whether INM PA levels increase in *pah1*1 *dgk1*1 cells (Fig. 6A). In the absence of choline, the PA sensor displayed predominantly a soluble nuclear distribution, both in wild-type and *pah1*1 *dgk1*1 cells (Fig. S6). Following choline addition, the sensor remained soluble in wild-type cells, while it strongly bound the deformed nuclear membranes in *pah1*1 *dgk1*1, indicative of a PA increase (Fig. S6). At late time points, the PA sensor became intranuclear again (Fig. S6, 120 min choline). The PA sensor-positive membranes surrounded an intranuclear reporter, confirming that they are indeed INM (Fig. 6B).

**Figure 6.**
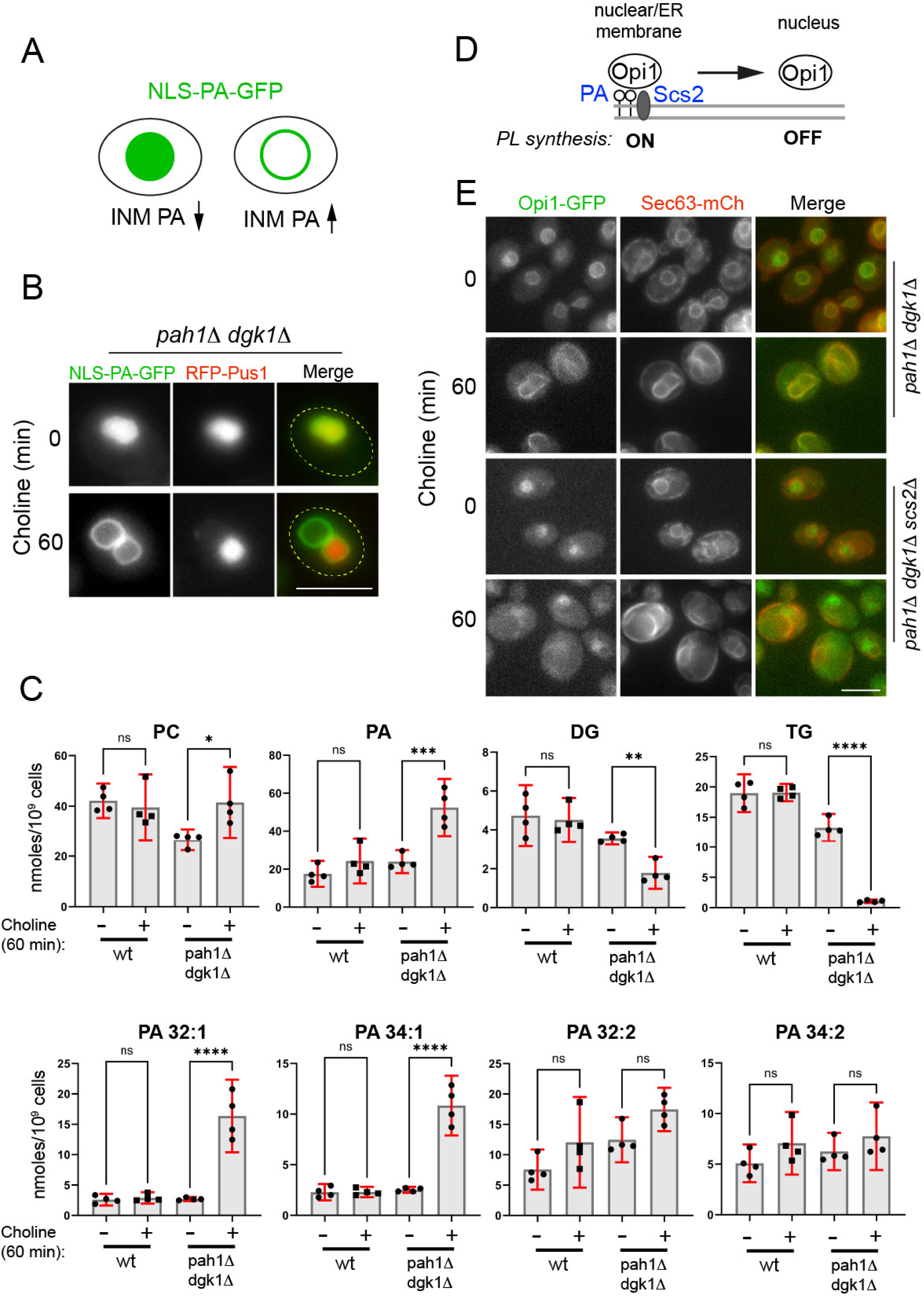
Pct1 is constitutively bound to PA-enriched INM and induces phospholipid synthesis. (A) Schematic of the behaviour of the PA biosensor at the INM. (B) *pah1*1 *dgk1*1 cells co-expressing the PA biosensor (NLS-PA-GFP) and RFP-Pus1 were supplemented with choline for the indicated times; the outlines of the cells are denoted. (C) *pah1*1 *dgk1*1 cells carrying two vectors (“*pah1*1 *dgk1*1”) or Pah1 and Dgk1 plasmids (“wt”) were grown in synthetic media, in the presence (1h) or absence of choline; cells were lysed and processed for lipidomics analysis as described in Materials and Methods; top row shows levels of cellular PC and PA (sum of 32:1, 32:2, 34:1 and 34:2 species), DG (sum of 32:1, 34:1, 34:2 and 36:1 species) and TG (sum of 42:0, 42:1, 42:2, 44:1, 44:2, 46:1, 46:2, 48:1, 48:2, 48:3, 50:2 and 50:3 species); lower row shows the denoted PA species; data are means +/- SD from four different transformants for each strain. Statistical significance was determined by one-way ANOVA with Šidák multiple-comparison test, or Welch ANOVA with Dunnett’s T3 test where variances were unequal. (D) Schematic of the regulation of Opi1. (E) The indicated strains expressing Opi1-GFP from its chromosomal locus and a plasmid-borne Sec63-mCh were supplemented with choline for the indicated times, and imaged; scale bars, 5 μm.

Next, to determine the lipid composition of the cells where Pct1 is constitutively membrane-bound, we used mass spectrometry. We compared the *pah1*1 *dgk1*1 strain bearing, either two empty vectors (“*pah1*1 *dgk1*1”) or complemented by both Dgk1 and Pah1 (“wild-type”) in the absence of choline or following its addition for 1 hour. In the wild-type strain, while individual PC species show modest changes (Fig. S7A), total PC levels were not significantly altered, consistent with previous reports that choline addition induces PC turnover and cells achieve homeostasis following one hour growth in choline-containing medium (Fig. 6C). In contrast, and in line with the observed membrane proliferation, total PC significantly increased in *pah1*1 *dgk1*1 following choline addition. PE levels also increased, while PS did not change and PI decreased (Fig. S7B). PA showed the largest increase of all phospholipid classes - more than 100%. The most significant increases were observed in the 32:1 and 34:1 species: in the absence of choline their levels were similar to those of the wild-type strain but both increased more than 200% after 1 hour in choline-supplemented media (Fig. 6C). Notably, following choline addition, PA increased even in the single *pah1*1 mutant, despite its already elevated basal PA levels (Fig. S7C). Concomitantly, DG levels in the *pah1*1 *dgk1*1 strain decreased by approximately 50%, consistent with its essential requirement as a precursor for PC synthesis via the Kennedy pathway (Fig. 6C). We further noticed that total *pah1*1 *dgk1*1 TG stores decreased dramatically after choline addition (Fig. 6C). Collectively, these data suggest that the failure to release Pct1 off PA-enriched INM in *pah1*1 *dgk1*1 leads to increased phospholipid biogenesis.

PA can also induce phospholipid synthesis via the Opi1 (Henry) circuit. Opi1 is a soluble transcriptional repressor of several phospholipid biosynthetic enzymes and is normally tethered at the perinuclear ER by both PA and the transmembrane protein Scs2 (Loewen et al., 2004; Loewen et al., 2003) (Fig. 6D); high levels of PA keep Opi1 membrane-bound and thus de-repress phospholipid synthesis via the methylation (Cho2-Opi3) pathway (Fig. 1A). To test whether the observed membrane proliferation in *pah1*1 *dgk1*1 requires a functional Opi1 circuit, we deleted Scs2, which causes Opi1 to remain in the nucleus and repress phospholipid synthesis even in the presence of high PA levels (Gaspar et al., 2017). Consistently, we found Opi1-GFP primarily at the perinuclear ER in *pah1*1 *dgk1*1 and intranuclear in *pah1*1 *dgk1*1 *scs2*1 (Fig. 6E). Choline addition led to similar membrane expansion in both strains, with Opi1-GFP remaining bound at the ER membrane in the presence of Scs2, and intranuclear in its absence (Fig. 6E). Therefore, while Opi1-mediated de-repression is likely to contribute to the elevated phospholipid levels, we propose that the observed membrane expansion in *pah1*1 *dgk1*1 is primarily driven by the Pct1-driven over-activation of the Kennedy pathway.

## Discussion

Prompted by the finding that the rate-limiting step of the Kennedy pathway occurs inside the nucleus across evolutionarily divergent models, we set out to investigate how this control point is regulated and how it is coordinated with downstream PC synthesis and its distribution across the ER-INM network. Our data support a model in which the Kennedy pathway functions as a homeostat for membrane growth: in wild-type cells, Pct1 cycles between a soluble nuclear pool and the INM, providing a reversible switch that matches the cellular demand for PC. In contrast, when Pct1 fails to dissociate from the INM, the Kennedy pathway is locked in an “ON” state, driving sustained CDP–choline production and, ultimately, excessive phospholipid synthesis and membrane biogenesis. We propose that this persistent Pct1 association can be triggered by elevated PA levels at the INM. PA is a potent activator of the Kennedy pathway because it can promote Pct1 membrane binding through two non-exclusive routes: (i) by increasing membrane packing defects and curvature elastic stress due to its cone-shaped form, and (ii) by strengthening electrostatic interactions via its anionic headgroup (Cornell, 2016). Thus, both bilayer physical properties and surface charge are likely combined to stabilize Pct1 on the INM, in agreement with *in vitro* data showing that lipid geometry and charge act synergistically to enhance CCTα membrane binding (Arnold and Cornell, 1996).

A likely cause of the acute choline-dependent PA increase in our *pah1*1 *dgk1*1 model is that PA production continues during choline addition, while its major route of removal, Pah1-mediated dephosphorylation to DG, is blocked. Consistent with this view, a *pah1*1 single mutant, which already exhibits elevated PA in the absence of choline, shows a further PA increase upon choline supplementation. The accompanying decrease in DG after choline addition is also consistent with the idea that the Kennedy pathway drives membrane proliferation by consuming DG for PC synthesis. However, this observation raises an important question: in the absence of Pah1, what is the source of DG to support continued phospholipid production? One possibility is that DG is supplied by neutral lipid mobilization. In line with this, TG levels in *pah1*1 *dgk1*1 drop sharply following choline supplementation, consistent with enhanced TG lipolysis providing DG for phospholipid synthesis. However, because basal TG stores in *pah1*1 *dgk1*1 are already reduced to half relative to the wild type, TG mobilization alone may be insufficient to account for sustained membrane biogenesis, implying that additional DG-generating routes may contribute. Together, these observations suggest extensive re-wiring of the lipid network in order for cells to keep utilizing choline even when key downstream steps are blocked. Choline is available as a nutrient in the wild, and yeasts possess dedicated transport and salvage pathways to process it (Fernandez-Murray et al., 2013; Patton-Vogt, 2007), suggesting that its uptake and utilization has evolved as a priority for yeast physiology.

Elevated PA can also promote membrane biogenesis indirectly by sequestering the transcriptional repressor Opi1 at the ER, thereby de-repressing UAS_INO_ genes, including those of the methylation branch (Cho2 and Opi3) that make PC (Henry et al., 2012). It is therefore plausible that the methylation branch contributes to the phospholipid accumulation in *pah1*1 *dgk1*1 cells. However, two observations argue that it is not the primary driver in our system. First, membrane expansion is Pct1-dependent; second, membrane proliferation persists even when Opi1 is relocalized to the nucleus. We therefore favor a model in which the constitutive membrane binding of Pct1 drives excessive PC synthesis. This model is also consistent with the PC species profile: the Kennedy pathway contributes significantly to newly synthesized monounsaturated PC species (Boumann et al., 2003), which are the same species that increase in *pah1*1 *dgk1*1 strain after choline supplementation; however, as alternative DG sources and post-synthetic acyl-chain remodelling could alter the PC profile during the choline feed, this interpretation should be considered with caution.

PA lies at the crossroads of multiple lipid metabolic pathways and consequently its levels in wild-type cells normally remain low. Thus, the sustained Pct1 activation described here is unlikely to occur under unperturbed growth. However, if PA levels rise briefly after choline uptake, and this surge exceeds the capacity of Pah1 to convert PA to DG, PA could stabilize Pct1 at the INM to increase nuclear CDP–choline production. Therefore, PA-mediated binding of Pct1 may serve as a rapid homeostat to coordinate nuclear CDP-choline with DG production by Pah1 until PC supply is restored. Notably, Pct1 is one of the few lipid metabolic enzymes whose expression does not appear to be regulated by PA at the transcriptional level (Jesch et al., 2006; Jesch et al., 2005). Therefore, the Kennedy pathway may be primarily regulated by a PA-mediated biochemical mechanism, as opposed to the methylation branch, which is primarily adjusted by PA through transcriptional feedback.

Our anchor-away experiments suggest that Pct1 inactivation is insensitive to where Kennedy-derived PC is produced. Our data are consistent with the idea that redistribution of PC is sufficiently rapid that it does not limit Pct1 inactivation at the time resolution of our assay. After choline supplementation, cells must import choline, phosphorylate it, convert it to CDP-choline and finally condense it with DG by Cpt1/Ept1 at the ER, before PC levels and curvature stress at the INM change. If any of these reactions proceed more slowly than PC bulk movement, the distance between PC production and sensing will have little impact on Pct1 translocation. Membrane contact sites are likely to accelerate lipid redistribution; given that 45% of the plasma membrane in yeast is closely apposed to the ER (Manford et al., 2012; West et al., 2011), we speculate that PC made by Pil1-anchored Cpt1 will rapidly enter the ER network. While the mechanisms of ER to INM lipid flow remain poorly defined, the ER membrane has a relatively disordered structure (Monje-Galvan and Klauda, 2015) and is continuous with the INM, allowing rapid phospholipid exchange between the nucleus and the rest of the ER. Consistent with dynamic equilibration of Kennedy-derived PC across the nuclear-ER membrane network, choline triggers a rapid change of nuclear shape, which requires Cpt1 and correlates with Pct1 translocation kinetics. An analogous robustness to altered subcellular sites of PE and PC synthesis has recently been reported (John Peter et al., 2022).

Pct1/CCTα do not detect PC directly but rather respond to the structural consequences of PC deficiency, which translate into membrane packing defects. The cytoplasmic endomembrane system is rich in membranes with complex shapes and charges and which respond to multiple stimuli other than PC deficiency (Bigay and Antonny, 2012). In contrast, the INM consists predominantly of a large flat membrane sheet making it a sensitive platform for sensing PC sufficiency and curvature stress. This could make the INM the ideal compartment for monitoring PC homeostasis across the ER-nuclear membrane continuum.

## Materials and methods

### Yeast strains and plasmids

Yeast strains are listed in Table S1 and were derived from BY4741, BY4742 or W303. Epitope tagging was performed either by (i) chromosomal integration at the endogenous locus or (ii) from centromeric (CEN/ARS) plasmids. Gene deletions and chromosomally integrated epitope tags were generated using a one-step PCR-based method (Janke et al., 2004; Longtine et al., 1998) and verified by PCR of genomic DNA. Plasmids are listed in Table S2. For plasmid-based tagging, epitope-tagged fusions and amino acid mutants were constructed by standard cloning procedures and PCR-mediated mutagenesis. All Pct1, Ept1 and Cpt1 constructs contained an in-frame BamHI restriction site immediately after the start codon, or before the stop codon, to enable insertion of tag-encoding fragments at the N- or C-terminus, respectively.

### Media and growth conditions

Cells were transformed using the lithium acetate method (Gietz and Woods, 2002). Cells were cultured in YEPD medium (1% bacto yeast extract, 2% peptone, 2% glucose) or in synthetic medium (SC) containing 0.17% yeast nitrogen base (Difco, BD), 0.5% ammonium sulfate, and amino acid drop-out containing 60 mg/L leucine, 55 mg/L adenine, 55 mg/L uracil, 55 mg/L tyrosine, 20 mg/L of arginine, 10 mg/L histidine, 60 mg/L isoleucine, 40 mg/L lysine, 60 mg/L phenylalanine, 50 mg/L threronine, 10 mg/L methionine, 40 mg/L tryptophan, lacking the constituent(s) required for plasmid selection. Unless otherwise specified, cells were assayed at the exponential phase of growth. For this, cells were pre-cultured the day before the experiment, diluted to an appropriate OD_600_ nm according on the doubling time of each strain, and then grown for approximately 15h to a final OD_600_ nm between 0.3 to 0.6. Yeast cultures were grown at shaking incubators (220 rpm and 30°C) unless otherwise specified. Choline was added at 1mM final concentration in the form of choline chloride (C-7527, Sigma) from a 100x stock. For dot spot growth assays, log phase cells from liquid cultures were diluted to OD600nm 0.2. Serial dilutions (5x) were spotted on the appropriate plates and incubated at 30°C for 1 to 3 days.

### Fluorescence microscopy

Cells were cultivated at 30°C in synthetic medium. Aliquots were harvested at the specified timepoints and promptly subjected to live imaging at room temperature. Our imaging setup comprised two microscopes. For the majority of experiments, we utilized a Zeiss AxioImager epifluorescence upright microscope equipped with a 100× Plan-Apochromatic 1.4 numerical aperture (NA) objective lens (Carl Zeiss, Jena, Germany). Image acquisition was conducted using a high-sensitivity fluorescence imaging large chip sCMOS mono camera (ORCA Flash 4, version 2; Hamamatsu, Hamamatsu, Japan). Zeiss ZEN blue software was employed to save raw image files, which were then exported to Fiji (ImageJ) and Adobe Photoshop. When specified, images were obtained using a Zeiss Axioplan epifluorescence microscope and a 100× Plan-Apochromatic 1.4 NA objective lens, connected to a Hamamatsu ORCA R2 charge-coupled device camera. Raw files were saved using Simple PCI6 software (Hamamatsu) and later exported to Fiji (ImageJ) and Adobe Photoshop.

All microscopy images were captures in a blinded manner. Quantitative analysis was carried out on fields derived from independent experiments.

### Pct1 nuclear distribution analysis

The quality and noise in fluorescent microscope images necessitated several pre-processing steps before graphically representing PCT1’s position. Initially, we created two separate images: one of PCT1 and another of PCT1 merged with a membrane reporter (Sur4-GFP or Sec63-GFP). These images were then aligned to form a composite montage, typically in a 5×1 arrangement. To enhance and clarify the signal, we applied the “subtract background” option using a rolling ball method with a 50-pixel radius. Subsequently, convolution kernels were applied, followed by four additional rounds of background subtraction using a 25-pixel rolling ball method, further refining the graph’s shape. Next, a line measuring 4.03 μm was drawn across the nucleus, defined by the overlap of Pct1 and the ER reporter signals. Measurements from this line were transferred to the PCT1-only channel. Signal intensity along a measurement line was normalized to the maximum value, squared to enhance profile shape, and values below 25% of the maximum were set to zero. Slopes were then calculated over a window of 4 to 10 points - consistent within an experiment - to determine whether the trace was increasing (coded as 1) or decreasing (coded as 0). The number of directional changes was then counted: profiles with more than two changes were classified as “Ring” (PCT1 membrane attachment), while those with two or fewer changes were classified as “Translocated” (nuclear accumulation).

### Propargyl choline click chemistry

Yeast cultures were divided into 2 ml tubes and rapamycin was added at a final concentration of 1 µg/ml to the anchored samples for 1 hour. Propargylcholine (Cayman Chemical) and choline (as a control) were added at 1 mM final concentration. Cells were collected at various time points between 0 and 30 min and fixed at room temperature while on a rotor by adding 37% formaldehyde (Sigma F-1635) at a final concentration of 3.7%. The cells were then spun at 4000 rpm for 4 minutes, the solution was removed, and the cells were placed in PBS/Sorbitol (0.1M PBS/1.2M Sorbitol) for 5 minutes. Afterward, the cells were spun again for 4 minutes at 4000 rpm and incubated for 5 minutes in 0.1 M Tris pH 8.5. Meanwhile, a click chemistry solution was prepared. Initially, ascorbic acid (Sigma 33034) was dissolved in 1M Tris pH 8.5 to a concentration of 0.3M. This solution was then added to CuSO4 (Fisher Scientific C/8520/50) to reach a final concentration of 1 mM and incubated at room temperature for 10 minutes. Subsequently, AlexaFluor405-Azide (4 µM), dissolved in 0.1M Tris pH 8.5, was added to achieve a final concentration of 1 µM. As a final step, the cells suspended in 0.1M Tris pH 8.5 were added to the solution, mixed, and incubated in darkness for 30 minutes. After this incubation, the cells were washed once with 0.1M Tris pH 8.5 before imaging.

### Microfluidics device fabrication

Microfluidic devices were fabricated by first making a master mould, using a soft photolithographic process through a previously established protocol (Czekalska et al., 2021; Hakala et al., 2021; Kartanas et al., 2021). In brief, a silicon wafer was placed under vacuum and 5µm photoresist (SU-8 3005, MicroChem) was spin-coated onto the wafer. This was then soft baked at 95°C for 3 min. Following this initial bake, the mask (the design of which was made on AutoCAD) was placed onto the wafer. This was subsequently exposed to UV light in order to induce polymerisation. The wafer was then baked a second time at 95 °C for 10 min. The master was then developed using propylene glycol methyl ether acetate (PGMEA, Sigma-Aldrich).

Following master fabrication, a device was prepared. This was achieved by adding the elastomer polydimethylsiloxane (PDMS) with curing agent (Sylgard 184, DowCorning, Midland, MI) at a ratio of 10:1. This was then placed under vacuum to remove any air bubbles for 1.5 hrs. The device was then incubated at 65 °C for 3 hours in order to induce curing. Following the curing process, the PDMS was cut out of the master and holes were punched, thus forming inlets and outlets. Finally, the cut PDMS slab was bound to a glass slide using a plasma bonder (Diener Electronic, Ebhausen, Germany).

### Microfluidics-based experiments

To perform the microfluidic experiments, we first had to fill the devices with cell solutions. Initially, plastic syringes were filled with the required cell solution. The syringe was attached to a needle which in turn was attached to plastic tubing. The tubing was placed in an inlet of the device, and the cell solution was injected into the microfluidic device (Toprakcioglu and Knowles, 2021). Once enough cells were trapped within the channels of the device, the tubing was removed from the inlet. A different syringe (e.g. a control syringe or one containing choline) was prepared using the same approach as described above and the tubing was inserted into the inlets of the microfluidic device. The device was placed onto the fluorescent microscope and image acquisition was initiated. The solution within the syringes was pumped through the microfluidic channels using syringe pumps. The flow rates within the microchannels were controlled using neMESYS syringe pumps (Cetoni).

### Immunoblotting

10 OD_600_ nm of yeast cells were pelleted, washed with water and resuspended in 160 μl of SDS-sample buffer with 0.5 mm diameter glass beads (BioSpec Products). Cell suspensions were lysed by two rounds of boiling for 2 min followed and vortexing for 30 s. Lysates were centrifuged at 13,000 rpm for 15 min, and the supernatants analyzed by western blot. Signals were developed by ECL (GE Healthcare). The anti-GFP antibody was from Abcam (ab6556).

### Mass spectrometry lipidomics

#### Cell growth and lysis

Exponentially growing *pah1*1 *dgk1*1 cells, carrying a chromosomally integrated *PCT1*-mCh fusion and either empty vectors, or *PAH1*/*DGK1* plasmids, were grown in synthetic medium (-Ura -Leu) with (1h) or without choline. Strain phenotypes were verified by microscopy prior harvesting. Cells were then pelleted, washed once with water, and snap-frozen in liquid nitrogen. Lipids were extracted from cell pellets as previously described (Folch et al., 1957), with minor modifications. Briefly, cell pellets were resuspended in a solution consisting of 650 μl chloroform, 250 μl methanol and 100 μl methanol containing the internal standard (described below). The cells were lysed with 100 mg 0.5 mm diameter glass beads (BioSpec Products) using a Precellys-24 homogenizer (Bertin) at 4°C and the following settings: 6500 rpm for 30 seconds, rest for 60 seconds, then repeating this cycle 5 times. Samples were sonicated with five short pulses and one minute on ice in between to prevent overheating. 400 μl of water was then added to the homogenates before the samples were vortexed for 30 sec and centrifuged for at 13,200 rpm at 4°C. The organic layer was then collected in a 2 mL amber glass vial (Agilent Technologies) and air-dried overnight in a fume hood.

#### Lipid analysis

For LC-MS analysis, the samples were reconstituted in 100 µL of 2: 1: 1 (propan-2-ol, acetonitrile and water, respectively) then thoroughly vortexed. The reconstituted sample was transferred into a 250 μL low-volume vial insert inside a 2 mL amber glass auto-sample vial ready for liquid chromatography with mass spectrometry detection (LC-MS) analysis. Full chromatographic separation of intact lipids was achieved using Waters Acquity H-Class HPLC System (Waters) with the injection of 10 µL onto a Waters Acquity Premier UPLC® CSH C18 column; 1.7 µm, I.D. 2.1 mm X 50 mm, maintained at 55 degrees Celsius. Mobile phase A was 6:4, acetonitrile and water with 10 mM ammonium formate. Mobile phase B was 9:1, propan-2-ol and acetonitrile with 10 mM ammonium formate. The flow was maintained at 500 µL per minute through the following gradient: 0.00 minutes 40% mobile phase B; 1.5 minutes 40% mobile phase B; 8.00 minutes 99% mobile phase B; 10.00 minutes 99% mobile phase B; 10.10 minutes 40% mobile phase B; 12.00 minutes 40% mobile phase B. The sample injection needle was washed using 9:1, propan-2-ol and acetonitrile [strong wash] and 2: 1: 1 (propan-2-ol, acetonitrile and water) [weak wash]. The mass spectrometer used was the Thermo Scientific Q-Exactive Orbitrap with a heated electrospray ionisation source (Thermo Fisher Scientific). The mass spectrometer was calibrated immediately before sample analysis using positive and negative ionisation calibration solution (recommended by Thermo Scientific). Additionally, the mass spectrometer scan rate was set at 4 *Hz*, giving a resolution of 35,000 (at 200 *m/z*) with a full-scan range of *m/z* 120 to 1,800 with continuous switching between positive and negative mode. Samples were analysed in groups of ten samples flanked by solvent blank samples, internal standard blank samples and quality control samples (Commercially available blank human serum was purchased from BioIVT (Royston, Hertfordshire, UK; order number: HUMANSRMPNN)).

The data processing was done using Thermo Scientific Xcalibur (Version 4.1.31.9) in the Quan browser. The integration of the extracted ion chromatogram peaks for each target lipid species and the stable isotope labelled internal standards at their expected retention time. The area ratio response of the target lipid to the corresponding internal standard were converted into nmole results normalised to cell count and subjected to blank correction and comprehensive quality checking before further statistical analysis.

#### Lipid Internal standard

Stable isotope-labelled internal standards purchased from Sigma Aldrich (Haverhill, Suffolk, UK) include: N-palmitoyl-d31-D-erythro-sphingosine (abbreviated to IS_Cer_16:0-d31); order number: 868516P, 1-palmitoyl-d31-2-oleoyl-sn-glycero-3-phosphate (abbreviated to IS_PA_34:1-d31); order number: 860453P, 1-palmitoyl-d31-2-oleoyl-sn-glycero-3-phosphocholine (abbreviated to IS_PC_34:1-d31); order number: 860399P, 1-palmitoyl-d31-2-oleoyl-sn-glycero-3-phosphoethanolamine (abbreviated to IS_PE_34:1-d31); order number: 860374P, 1-palmitoyl-d31-2-oleoyl-sn-glycero-3-[phospho-rac-(1-glycerol)] (abbreviated to IS_PG_34:1-d31); order number: 860384P, 1-palmitoyl-d31-2-oleoyl-sn-glycero-3-phosphoinositol (abbreviated to IS_PI_34:1-d31); order number: 860042P, 1,2-dimyristoyl-d54-sn-glycero-3-[phospho-L-serine] (abbreviated to IS_PS_28:0-d54); order number: 860401P, N-palmitoyl-d31-D-erythro-sphingosylphosphorylcholine (abbreviated to IS_SM_34:1-d31); order number: 868584P. Stable isotope-labelled internal standards purchased from QMX Laboratories Ltd. (QMX Laboratories Ltd., Thaxted, Essex, UK) include: Heptadecanoic-d33 acid (abbreviated to IS_FA_17:0-d33); order number: D-5261, N-tetradecylphosphocholine-d42 (abbreviated to IS_LPC_14:0-d42); order number: D-5885, Glyceryl tri(pentadecanoate-d29) (abbreviated to IS_TG_45:0-d87); order number: D-5265, Butyryl-d7-L-carnitine (abbreviated to IS_Car_4:0-d7); order number: D-7761, Hexadecanoyl-L-carnitine-d3 (abbreviated to IS_Car_16:0-d3); order number: D-6646. The lipid stable isotope-labelled internal standard was prepared by dissolving each of the individual lipid standards into chloroform: methanol (1:1) solution to produce a 1 mM primary stock solution. From each of these stock solutions, 1 mL was transferred into a volumetric flask and diluted with methanol to reach a final working solution concentration of 5 µM in methanol of IS_Cer_16:0-d31, IS_FA_17:0-d33, IS_LPC_14:0-d42, IS_PA_34:1-d31, IS_PC_34:1-d31, IS_PE_34:1-d31, IS_PG_34:1-d31, IS_PI_34:1-d31, IS_PS_28:0-d54, IS_SM_34:1-d31, IS_TG_45:0-d87.

## Statistical analysis

Graphs and statistical analyses were performed using GraphPad Prism. Data are presented as mean ± standard deviation (SD). Variance equality was assessed using the Brown–Forsythe and Bartlett’s tests. When both tests indicated equal variances, ordinary one-way ANOVA followed by Šidák correction for multiple comparisons was used. When either Brown–Forsythe or Bartlett’s test indicated unequal variances, Welch ANOVA followed by Dunnett’s T3 correction for multiple comparisons was applied. p values were coded as follows: *P < 0.05, **P < 0.01, ***P < 0.001, and ****P <0.0001.

## Acknowledgments

We thank Kasparas Petkevicius for comments on the manuscript; Matthew Gratian and Mark Bowen for help with microscopy; and Alwin Köhler for plasmids. This work was supported by: Biotechnology and Biological Sciences Research Council grant [BB/T005610/1] (SS); Wellcome Trust grant 108415/Z/15/Z to the Cambridge Institute for Medical Research; Wellcome Trust grant WT 219417 (DBS); Wellcome Trust grant 226800/Z/22/Z (AK); the Ron Thomson Research Fellowship in Alzheimer’s Disease Pembroke College Cambridge (ZT); and the European Research Council under the European Union’s Horizon 2020 research and innovation program through the ERC grant DiProPhys (agreement ID 101001615) and the Francis and Augustus Newman Foundation (TPJK).

## Supplementary figure legends

**Figure S1.**
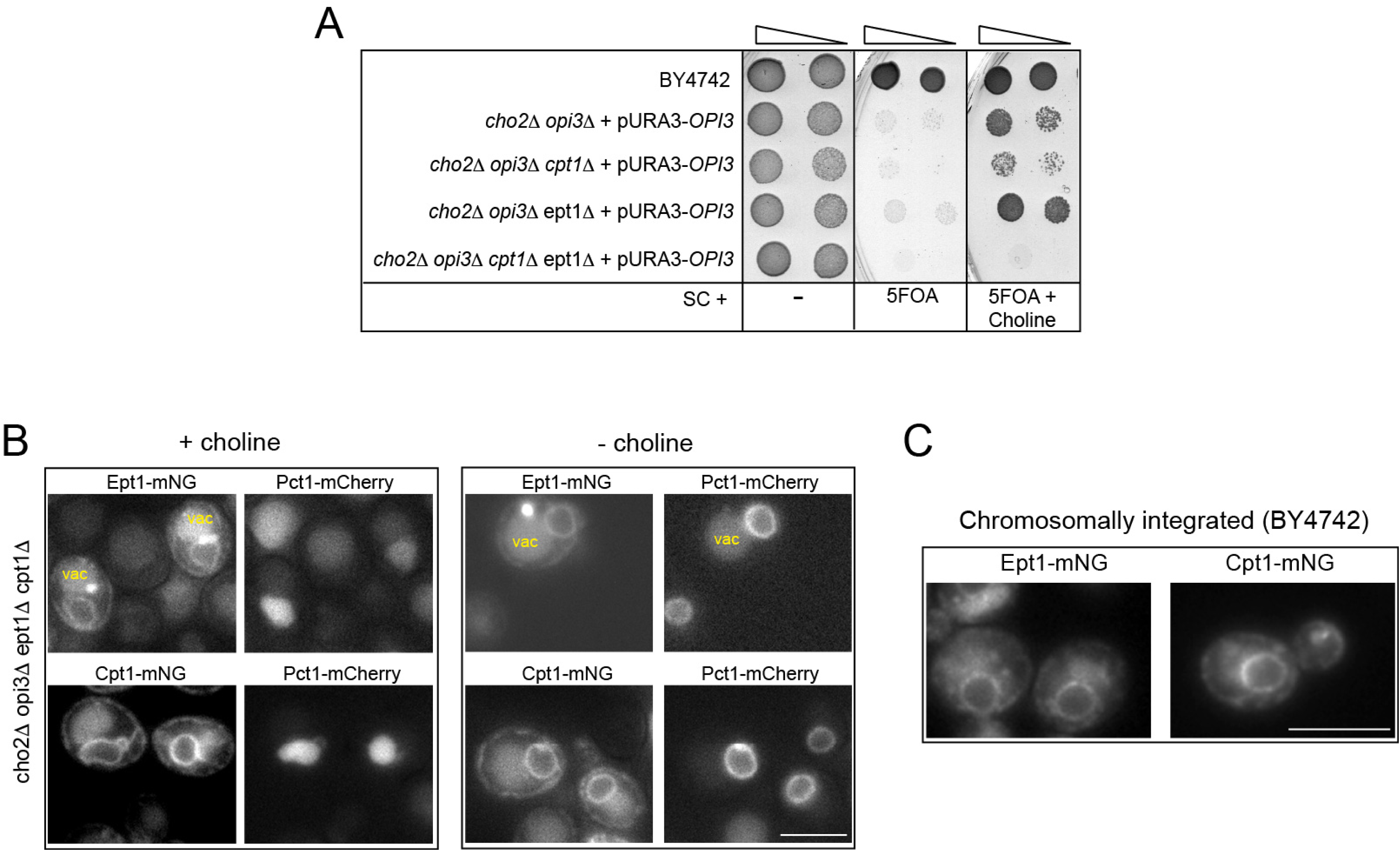
Distribution and functional requirement of Ept1 and Cpt1. (A) The denoted mutants disrupting the methylation or the Kennedy pathways, were spotted on plates in the absence or presence of 5-FOA to counter select the pURA3-OPI3 plasmid. Plates were supplemented with or without 1mM choline. (B) Ept1-mNG Cpt1-mNG were expressed in the ΔPC strain expressing Pct1-mCh in the presence of 1mM choline, or following 8h choline depletion. In some cells, Ept1-mNG localizes also to a perivacuolar dot, not seen with mNG-Ept1; vac: vacuole. (C) Cells expressing the indicated fusions were visualized as above. Scale bars, 5 μm.

**Figure S2.**
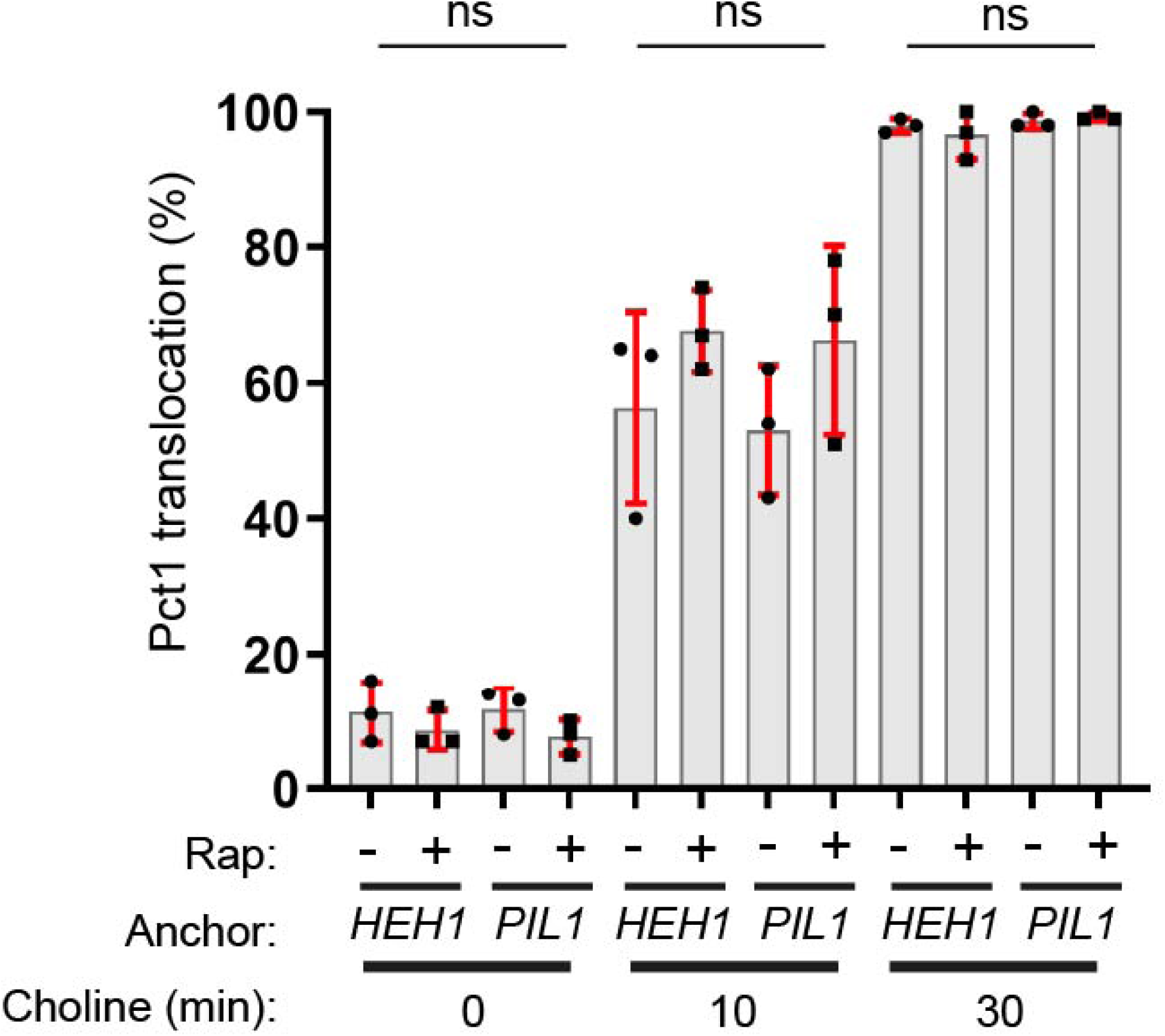
Pct1 nuclear translocation is independent of the site of PC synthesis. Quantification of Pct1 translocation shown in *CHO2 OPI3* cells; the anchor, the presence of rapamycin and the duration of choline supplementation, are denoted; data are means from three experiments +/- SD and analysed using a one-way ANOVA with Sidak’s post hoc test; ns, p≥ 0.05.

**Figure S3.**
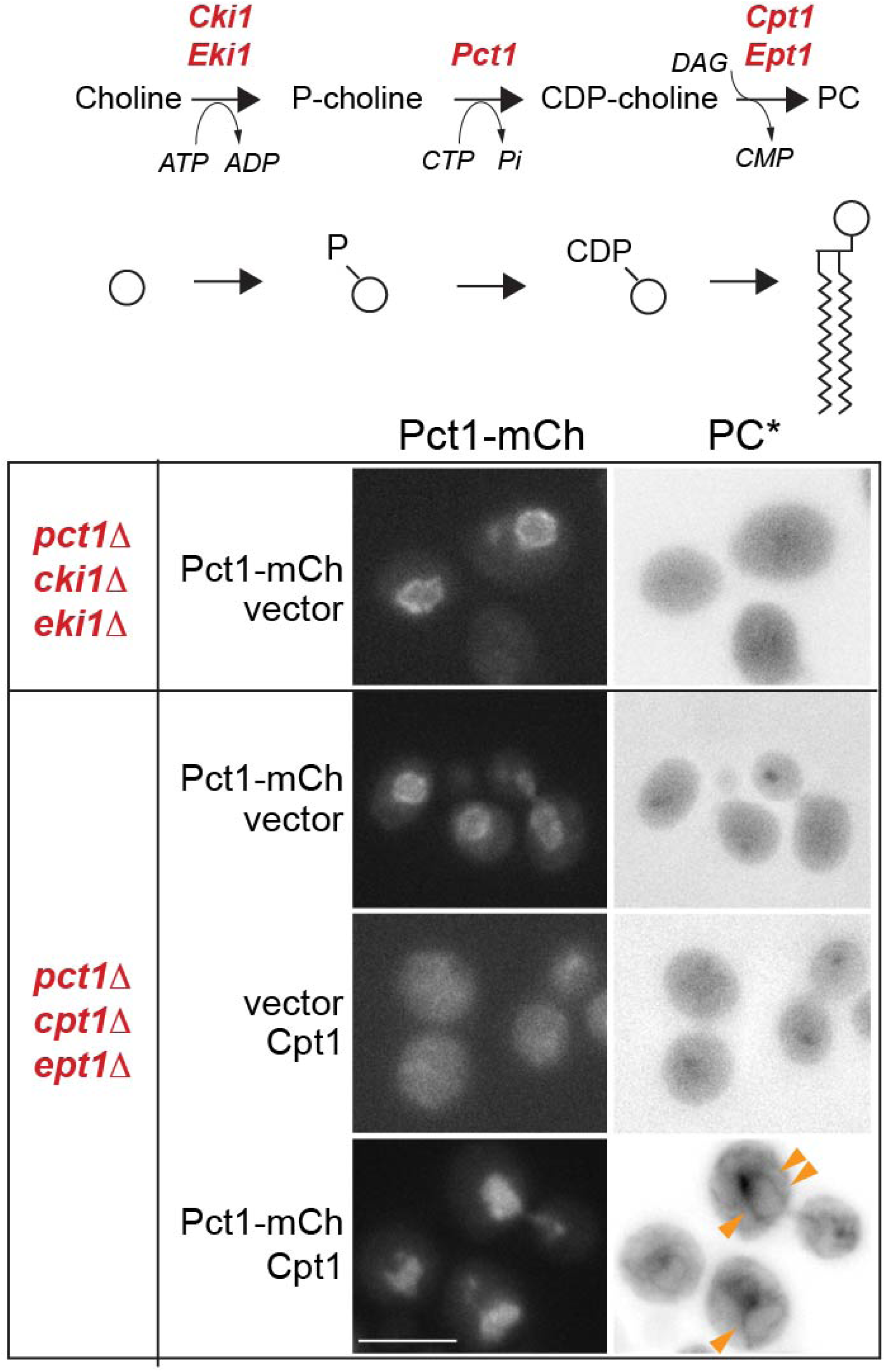
Membrane labelling of Pr-Cho requires a functional Kennedy pathway. (A) Schematic of the Kennedy pathway and the relevant enzymes. (B) Cultures of the denoted strains expressing the indicated plasmids, were supplemented with propargyl-choline for 30 min, fixed, and phosphatidyl-propargyl-choline (PC*) was visualized by click-chemistry. PC* micrographs are shown as inverted grey-scale images; arrows indicate nuclear membranes containing PC*. Scale bars, 5 μm.

**Figure S4.**
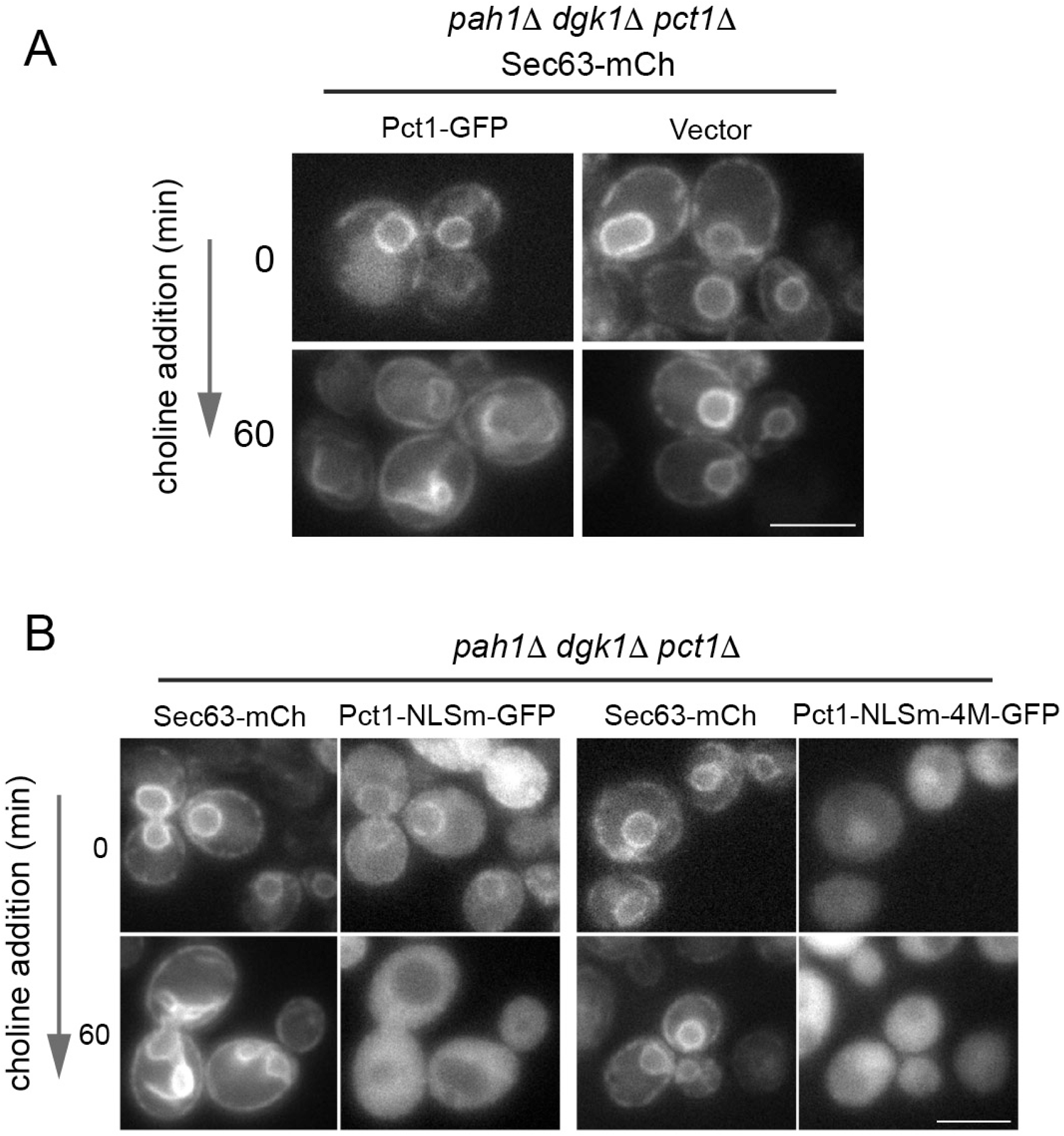
(A) Pct1 is required for membrane expansion in the presence of choline. *pah1*1 *dgk1*1 *pct1*1 cells, co-expressing Sec63-mCh and either Pct1-GFP or a vector, were supplemented with choline and imaged at the indicated time points. (B) Cytoplasmic Pct1 requires its membrane binding helix to drive membrane expansion. *pah1*1 *dgk1*1 *pct1*1 cells, co-expressing Sec63-mCh and Pct1-NLSm-GFP or a Pct1-NLSm-4M-GFP, were supplemented with choline and imaged at the indicated time points. Scale bars, 5 μm.

**Figure S5.**
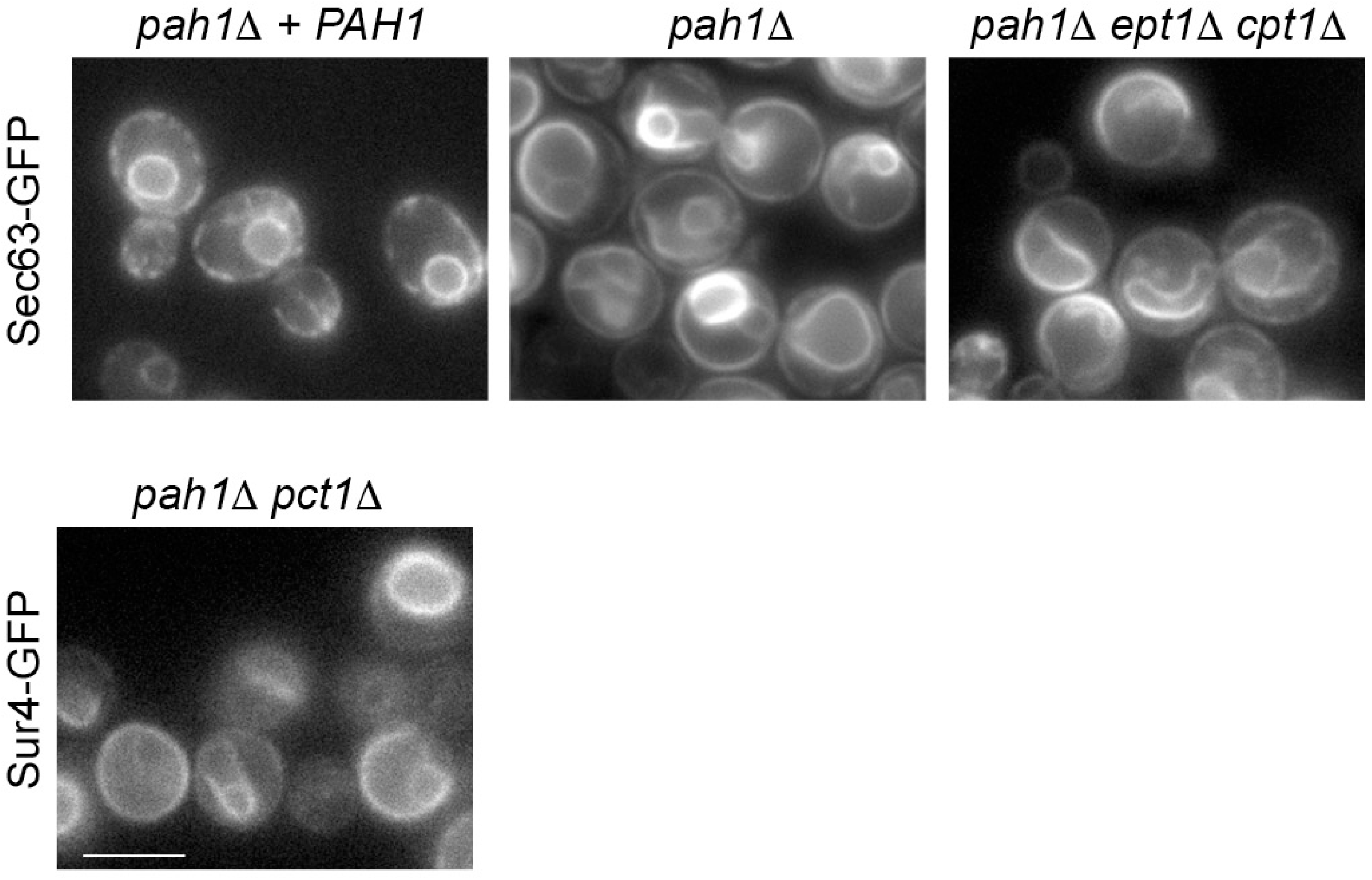
Disruption of the Kennedy pathway does not restore membrane homeostasis in *pah1*1 cells. The denoted strains expressing an ER membrane reporter - Sec63-GFP or Sur4-GFP – were grown in synthetic media and the ER morphology was visualized. Scale bars, 5 μm.

**Figure S6.**
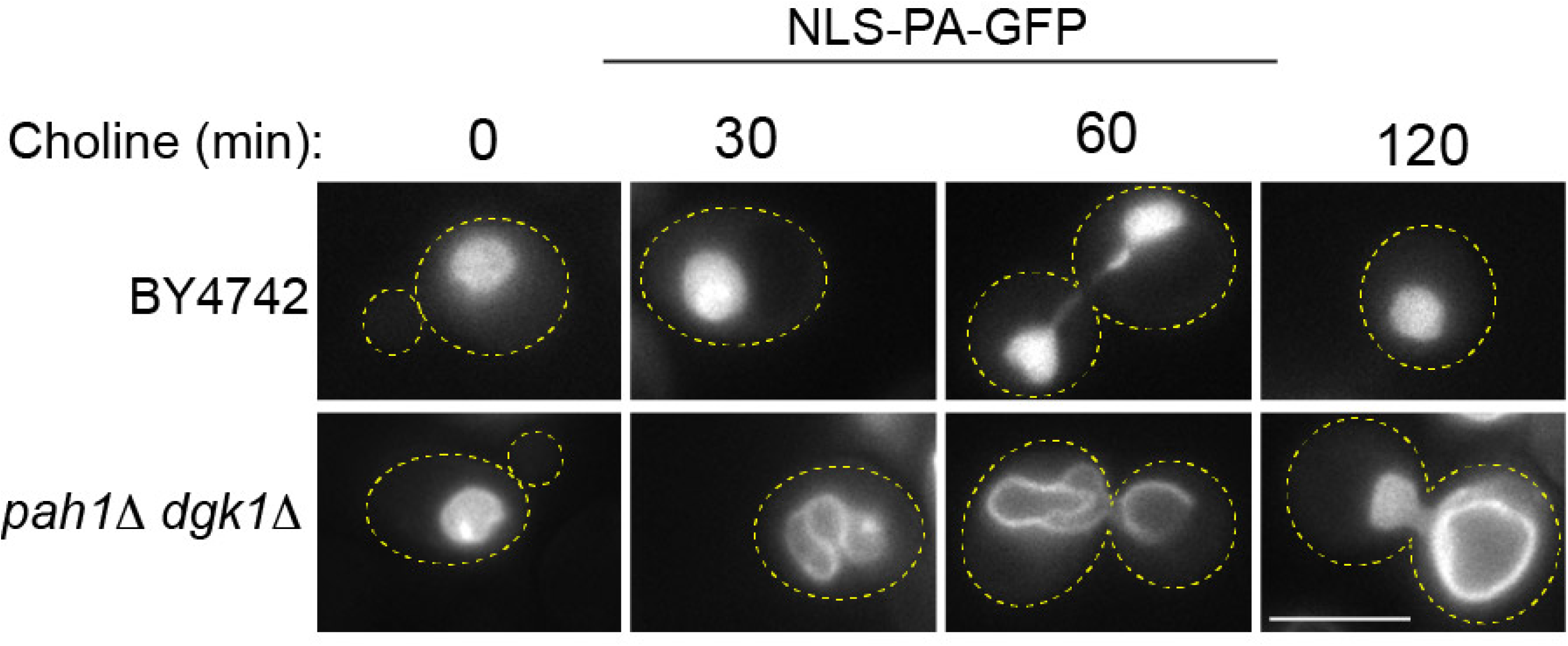
A PA biosensor binds the INM of *pah1*1 *dgk1*1 cells. Wild-type or *pah1*1 *dgk1*1 cells expressing the PA biosensor were supplemented with choline for the indicated times; the outline of cells is denoted. Scale bars, 5 μm.

**Figure S7.**
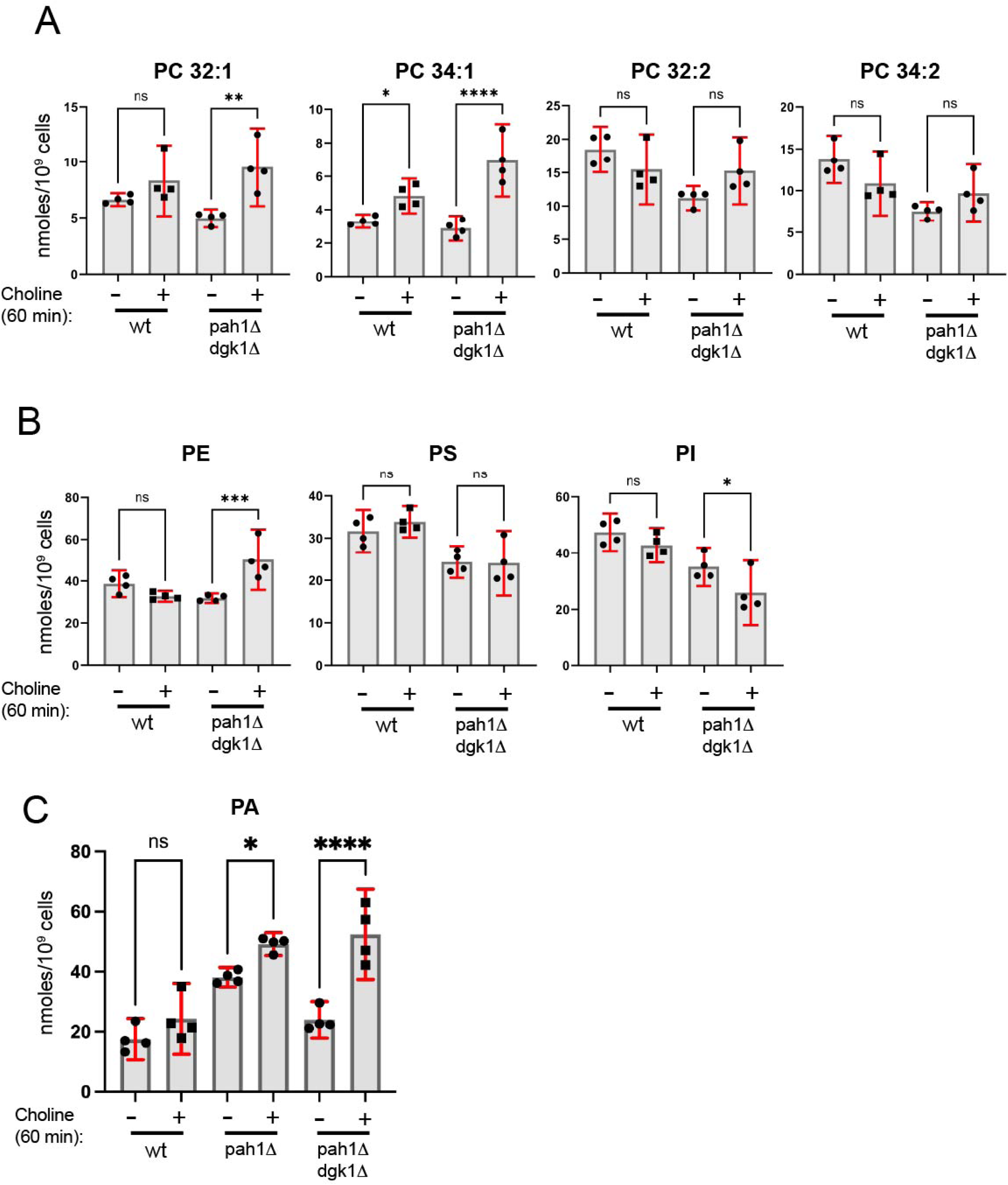
Quantification of the major PC species in (A) and cellular PE (sum of 32:1, 32:2, 34:1 and 34:2 species), PS (sum of 34:2 and 34:1 species) and PI (sum of 36:1, 34:1 and 34:2 species) in (B); *pah1*1 *dgk1*1 cells carrying two empty vectors (“*pah1*1 *dgk1*1”) or Pah1 and Dgk1 plasmids (“wt”) were grown in synthetic media, in the presence (1h) or absence of choline; cells were lysed and processed for lipidomics analysis as described in Materials and methods; data are means and SD’s from four different transformants for each strain; statistical significance was determined by one-way ANOVA with Šidák multiple-comparison test, or Welch ANOVA with Dunnett’s T3 correction when variances were unequal. (C) Quantification of cellular PA levels (sum of 32:1, 32:2, 34:1 and 34:2 species) in *pah1*1 *dgk1*1 cells carrying two vectors (“*pah1*1 *dgk1*1”), Dgk1 and an empty vector (“*pah1*1”), or Pah1 and Dgk1 plasmids (“wt”) in the presence (1h) or absence of choline. Note that these are the same samples as those shown in Fig. 6C but with the addition of the *pah1*1 strain.

---

**Table S1.**
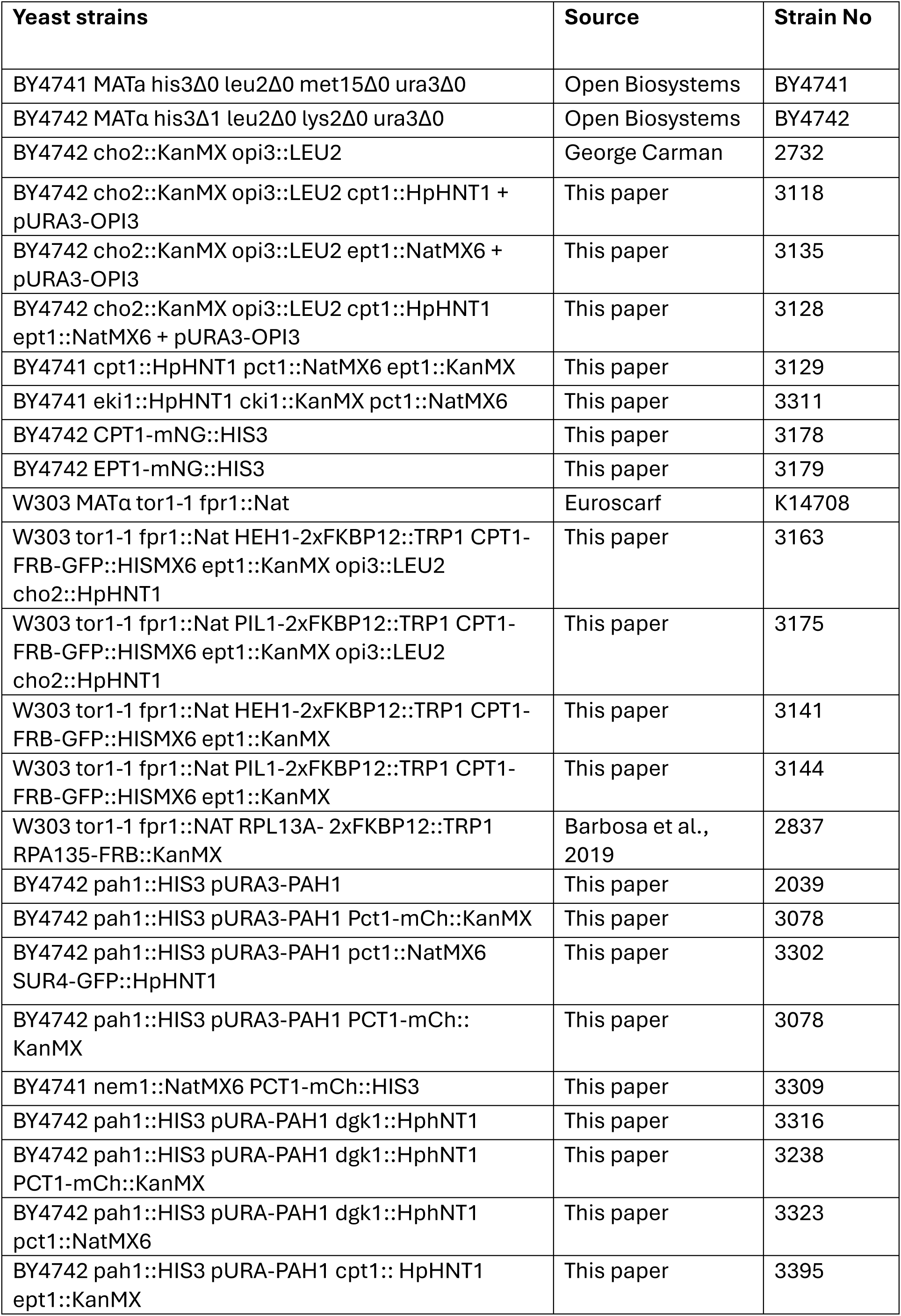
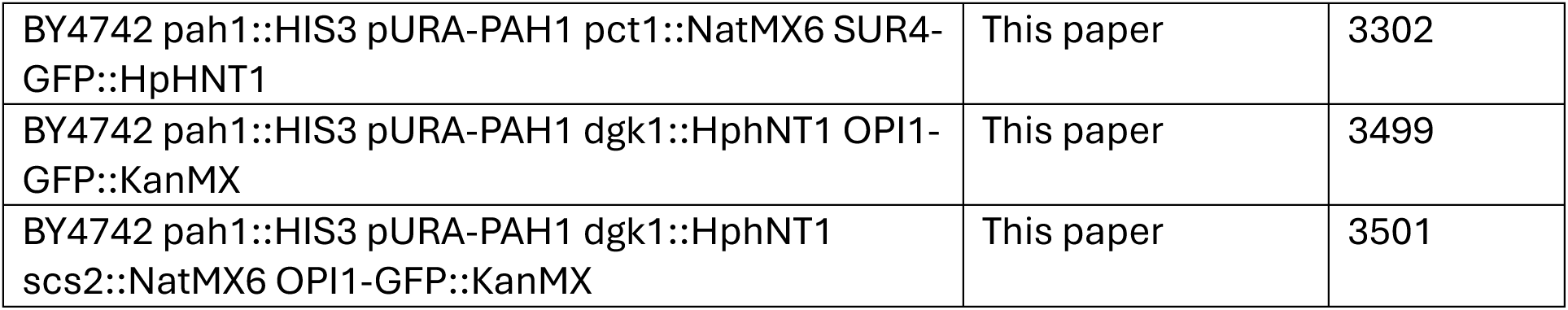
Yeast strains used in this study.

**Table S2.** Plasmids used in this study.

| Plasmid description | Source/reference | Name |
| --- | --- | --- |
| *EPT1* under control of *EPT1* promoter in *CEN/HIS3* vector | This paper | pRS313-*EPT1* |
| *CPT1* under control of *CPT1* promoter in *CEN/HIS3* vector | This paper | pRS313-*CPT1* |
| *mNG-EPT1* under control of *EPT1* promoter in *CEN/HIS3* vector | This paper | pRS313-*mNG-EPT1* |
| *EPT1-mNG* under control of *EPT1* promoter in *CEN/HIS3* vector | This paper | pRS313-*EPT1-mNG* |
| *mNG-CPT1* under control of *CPT1* promoter in *CEN/HIS3* vector | This paper | pRS313-*mNG-CPT1* |
| *CPT1-mNG* under control of *CPT1* promoter in *CEN/HIS3* vector | This paper | pRS313-*CPT1-mNG* |
| *PCT1-mCh* under control of *PCT1* promoter in *CEN/URA3* vector | This paper | YCplac33-*PCT1-mCh* |
| *PCT1-GFP* under control of *PCT1* promoter in *CEN/LEU2* vector | Haider et al., 2018 | YCplac111-*PCT1-GFP* |
| *SEC63-mCh* under control of *SEC63* promoter in *CEN/URA3* vector | Haider et al., 2018 | YCplac33-*SEC63-mCh* |
| *PAH1* under control of *PAH1* promoter in *CEN/LEU2* vector | O'Hara et al., 2006 | YCplac111-*PAH1* |
| *PAH1* under control of *PAH1* promoter in *CEN/URA3* vector | Han et al., 2008a | YCplac33-*PAH1* |
| *NEM1* under control of *NEM1* promoter in *CEN/URA3* vector | This paper | YCplac33-*NEM1* |
| *HEH2-GFP* under control of *NOP1* promoter in *CEN/LEU2* vector | Han et al., 2008 | YCplac111-*HEH2-GFP* |
| *PCT1-4M-GFP* under control of *PCT1* promoter in *CEN/LEU2* vector | This paper | YCplac111-*PCT1-4M-GFP* |
| *PCT1-NLSm-GFP* under control of *PCT1* promoter in *CEN/LEU2* vector | This paper | YCplac111-*PCT1-NLSm-GFP* |
| *PCT1-NLSm-4M-GFP* under control of *PCT1* promoter in *CEN/LEU2* vector | This paper | YCplac111-*PCT1-NLSm-4M-GFP* |
| *RFP-PUS1* under control of *NOP1* promoter in *CEN/LEU2* vector | This paper | YCplac111-*RFP-PUS1* |
| NLS-Opi1 Q2-mGFP: pRS316-CYC1prom-NUP60(1-24)-OPI1 Q2-mGFP | Romanauska and Köhler, 2018 | NLS-PA-GFP |
| *DGK1* under control of *DGK1* promoter in *CEN/LEU2* vector | This paper | YCplac111-*DGK1* |

## Notes

### Competing Interest Statement

The authors have declared no competing interest.

## References

1. Arnold, R.S., and R.B. Cornell. 1996. Lipid regulation of CTP: phosphocholine cytidylyltransferase: electrostatic, hydrophobic, and synergistic interactions of anionic phospholipids and diacylglycerol. Biochemistry. 35:9917–9924.

2. Bigay, J., and B. Antonny. 2012. Curvature, lipid packing, and electrostatics of membrane organelles: defining cellular territories in determining specificity. Dev Cell. 23:886–895.

3. Boumann, H.A., M.J. Damen, C. Versluis, A.J. Heck, B. de Kruijff, and A.I. de Kroon. 2003. The two biosynthetic routes leading to phosphatidylcholine in yeast produce different sets of molecular species. Evidence for lipid remodeling. Biochemistry. 42:3054–3059.

4. Boumann, H.A., B. de Kruijff, A.J. Heck, and A.I. de Kroon. 2004. The selective utilization of substrates in vivo by the phosphatidylethanolamine and phosphatidylcholine biosynthetic enzymes Ept1p and Cpt1p in yeast. FEBS Lett. 569:173–177.

5. Boumann, H.A., J. Gubbens, M.C. Koorengevel, C.S. Oh, C.E. Martin, A.J. Heck, J. Patton-Vogt, S.A. Henry, B. de Kruijff, and A.I. de Kroon. 2006. Depletion of phosphatidylcholine in yeast induces shortening and increased saturation of the lipid acyl chains: evidence for regulation of intrinsic membrane curvature in a eukaryote. Mol Biol Cell. 17:1006–1017.

6. Cornell, R.B. 2016. Membrane lipid compositional sensing by the inducible amphipathic helix of CCT. Biochim Biophys Acta. 1861:847–861.

7. Cornell, R.B. 2020. Membrane Lipids Assist Catalysis by CTP: Phosphocholine Cytidylyltransferase. J Mol Biol. 432:5023–5042.

8. Cornell, R.B., and I.C. Northwood. 2000. Regulation of CTP:phosphocholine cytidylyltransferase by amphitropism and relocalization. Trends Biochem Sci. 25:441–447.

9. Cornell, R.B., and N.D. Ridgway. 2015. CTP:phosphocholine cytidylyltransferase: Function, regulation, and structure of an amphitropic enzyme required for membrane biogenesis. Prog Lipid Res. 59:147–171.

10. Craddock, C.P., N. Adams, F.M. Bryant, S. Kurup, and P.J. Eastmond. 2015. PHOSPHATIDIC ACID PHOSPHOHYDROLASE Regulates Phosphatidylcholine Biosynthesis in Arabidopsis by Phosphatidic Acid-Mediated Activation of CTP:PHOSPHOCHOLINE CYTIDYLYLTRANSFERASE Activity. Plant Cell. 27:1251–1264.

11. Czekalska, M.A., A.M.J. Jacobs, Z. Toprakcioglu, L. Kong, K.N. Baumann, H. Gang, G. Zubaite, R. Ye, B. Mu, A. Levin, W.T.S. Huck, and T.P.J. Knowles. 2021. One-Step Generation of Multisomes from Lipid-Stabilized Double Emulsions. ACS Appl Mater Interfaces. 13:6739–6747.

12. Fagone, P., and S. Jackowski. 2013. Phosphatidylcholine and the CDP-choline cycle. Biochim Biophys Acta. 1831:523–532.

13. Fernandez-Murray, J.P., M.H. Ngo, and C.R. McMaster. 2013. Choline transport activity regulates phosphatidylcholine synthesis through choline transporter Hnm1 stability. J Biol Chem. 288:36106–36115.

14. Folch, J., M. Lees, and G.H. Sloane Stanley. 1957. A simple method for the isolation and purification of total lipides from animal tissues. J Biol Chem. 226:497–509.

15. Gaspar, M.L., Y.F. Chang, S.A. Jesch, M. Aregullin, and S.A. Henry. 2017. Interaction between repressor Opi1p and ER membrane protein Scs2p facilitates transit of phosphatidic acid from the ER to mitochondria and is essential for INO1 gene expression in the presence of choline. J Biol Chem. 292:18713–18728.

16. Gautier, R., D. Douguet, B. Antonny, and G. Drin. 2008. HELIQUEST: a web server to screen sequences with specific alpha-helical properties. Bioinformatics. 24:2101–2102.

17. Gietz, R.D., and R.A. Woods. 2002. Transformation of yeast by lithium acetate/single-stranded carrier DNA/polyethylene glycol method. Methods Enzymol. 350:87–96.

18. Haider, A., Y.C. Wei, K. Lim, A.D. Barbosa, C.H. Liu, U. Weber, M. Mlodzik, K. Oras, S. Collier, M.M. Hussain, L. Dong, S. Patel, A. Alvarez-Guaita, V. Saudek, B.J. Jenkins, A. Koulman, M.K. Dymond, R.C. Hardie, S. Siniossoglou, and D.B. Savage. 2018. PCYT1A Regulates Phosphatidylcholine Homeostasis from the Inner Nuclear Membrane in Response to Membrane Stored Curvature Elastic Stress. Dev Cell. 45:481–495 e488.

19. Hakala, T.A., E.V. Yates, P.K. Challa, Z. Toprakcioglu, K. Nadendla, D. Matak-Vinkovic, C.M. Dobson, R. Martinez, F. Corzana, T.P.J. Knowles, and G.J.L. Bernardes. 2021. Accelerating Reaction Rates of Biomolecules by Using Shear Stress in Artificial Capillary Systems. J Am Chem Soc. 143:16401–16410.

20. Han, G.S., L. O’Hara, G.M. Carman, and S. Siniossoglou. 2008a. An unconventional diacylglycerol kinase that regulates phospholipid synthesis and nuclear membrane growth. J Biol Chem. 283:20433–20442.

21. Han, G.S., L. O’Hara, S. Siniossoglou, and G.M. Carman. 2008b. Characterization of the yeast DGK1-encoded CTP-dependent diacylglycerol kinase. J Biol Chem. 283:20443–20453.

22. Han, G.S., W.I. Wu, and G.M. Carman. 2006. The Saccharomyces cerevisiae Lipin homolog is a Mg2+-dependent phosphatidate phosphatase enzyme. J Biol Chem. 281:9210–9218.

23. Haruki, H., J. Nishikawa, and U.K. Laemmli. 2008. The anchor-away technique: rapid, conditional establishment of yeast mutant phenotypes. Mol Cell. 31:925–932.

24. Henry, S.A., S.D. Kohlwein, and G.M. Carman. 2012. Metabolism and regulation of glycerolipids in the yeast Saccharomyces cerevisiae. Genetics. 190:317–349.

25. Hjelmstad, R.H., and R.M. Bell. 1991. sn-1,2-diacylglycerol choline- and ethanolaminephosphotransferases in Saccharomyces cerevisiae. Mixed micellar analysis of the CPT1 and EPT1 gene products. J Biol Chem. 266:4357–4365.

26. Iyoshi, S., J. Cheng, T. Tatematsu, S. Takatori, M. Taki, Y. Yamamoto, A. Salic, and T. Fujimoto. 2014. Asymmetrical distribution of choline phospholipids revealed by click chemistry and freeze-fracture electron microscopy. ACS Chem Biol. 9:2217–2222.

27. Janke, C., M.M. Magiera, N. Rathfelder, C. Taxis, S. Reber, H. Maekawa, A. Moreno-Borchart, G. Doenges, E. Schwob, E. Schiebel, and M. Knop. 2004. A versatile toolbox for PCR-based tagging of yeast genes: new fluorescent proteins, more markers and promoter substitution cassettes. Yeast. 21:947–962.

28. Jao, C.Y., M. Roth, R. Welti, and A. Salic. 2009. Metabolic labeling and direct imaging of choline phospholipids in vivo. Proc Natl Acad Sci U S A. 106:15332–15337.

29. Jesch, S.A., P. Liu, X. Zhao, M.T. Wells, and S.A. Henry. 2006. Multiple endoplasmic reticulum-to-nucleus signaling pathways coordinate phospholipid metabolism with gene expression by distinct mechanisms. J Biol Chem. 281:24070–24083.

30. Jesch, S.A., X. Zhao, M.T. Wells, and S.A. Henry. 2005. Genome-wide analysis reveals inositol, not choline, as the major effector of Ino2p-Ino4p and unfolded protein response target gene expression in yeast. J Biol Chem. 280:9106–9118.

31. John Peter, A.T., S.N.S. van Schie, N.J. Cheung, A.H. Michel, M. Peter, and B. Kornmann. 2022. Rewiring phospholipid biosynthesis reveals resilience to membrane perturbations and uncovers regulators of lipid homeostasis. EMBO J. 41:e109998.

32. Kartanas, T., A. Levin, Z. Toprakcioglu, T. Scheidt, T.A. Hakala, J. Charmet, and T.P.J. Knowles. 2021. Label-Free Protein Analysis Using Liquid Chromatography with Gravimetric Detection. Anal Chem. 93:2848–2853.

33. King, M.C., C.P. Lusk, and G. Blobel. 2006. Karyopherin-mediated import of integral inner nuclear membrane proteins. Nature. 442:1003–1007.

34. Kodaki, T., and S. Yamashita. 1989. Characterization of the methyltransferases in the yeast phosphatidylethanolamine methylation pathway by selective gene disruption. Eur J Biochem. 185:243–251.

35. Krahmer, N., Y. Guo, F. Wilfling, M. Hilger, S. Lingrell, K. Heger, H.W. Newman, M. Schmidt-Supprian, D.E. Vance, M. Mann, R.V. Farese, Jr., and T.C. Walther. 2011. Phosphatidylcholine synthesis for lipid droplet expansion is mediated by localized activation of CTP:phosphocholine cytidylyltransferase. Cell Metab. 14:504–515.

36. Loewen, C.J., M.L. Gaspar, S.A. Jesch, C. Delon, N.T. Ktistakis, S.A. Henry, and T.P. Levine. 2004. Phospholipid metabolism regulated by a transcription factor sensing phosphatidic acid. Science. 304:1644–1647.

37. Loewen, C.J., A. Roy, and T.P. Levine. 2003. A conserved ER targeting motif in three families of lipid binding proteins and in Opi1p binds VAP. EMBO J. 22:2025–2035.

38. Longtine, M.S., A. McKenzie, 3rd, D.J. Demarini, N.G. Shah, A. Wach, A. Brachat, P. Philippsen, and J.R. Pringle. 1998. Additional modules for versatile and economical PCR-based gene deletion and modification in Saccharomyces cerevisiae. Yeast. 14:953–961.

39. MacKinnon, M.A., A.J. Curwin, G.J. Gaspard, A.B. Suraci, J.P. Fernandez-Murray, and C.R. McMaster. 2009. The Kap60-Kap95 karyopherin complex directly regulates phosphatidylcholine synthesis. J Biol Chem. 284:7376–7384.

40. Manford, A.G., C.J. Stefan, H.L. Yuan, J.A. Macgurn, and S.D. Emr. 2012. ER-to-plasma membrane tethering proteins regulate cell signaling and ER morphology. Dev Cell. 23:1129–1140.

41. McMaster, C.R., and R.M. Bell. 1994. Phosphatidylcholine biosynthesis in Saccharomyces cerevisiae. Regulatory insights from studies employing null and chimeric sn-1,2-diacylglycerol choline- and ethanolaminephosphotransferases. J Biol Chem. 269:28010–28016.

42. Monje-Galvan, V.M., and J.B. Klauda. 2015. Modeling yeast organelle membranes and how lipid diversity influences bilayer properties. Biochemistry. 54:6852–6861.

43. Niu, Y., J.G. Pemberton, Y.J. Kim, and T. Balla. 2024. Phosphatidylserine enrichment in the nuclear membrane regulates key enzymes of phosphatidylcholine synthesis. EMBO J. 43:3414–3449.

44. O’Hara, L., Han, G.S., Peak-Chew, S., Grimsey, N., Carman, G.M., and S. Siniossoglou. 2006. Control of phospholipid synthesis by phosphorylation of the yeast lipin Pah1p/Smp2p Mg2+-dependent phosphatidate phosphatase. J Biol Chem. 281:34537–34548.

45. Patton-Vogt, J. 2007. Transport and metabolism of glycerophosphodiesters produced through phospholipid deacylation. Biochim Biophys Acta. 1771:337–342.

46. Ridgway, N.D., and D.E. Vance. 1987. Purification of phosphatidylethanolamine N-methyltransferase from rat liver. J Biol Chem. 262:17231–17239.

47. Romanauska, A., and A. Kohler. 2018. The Inner Nuclear Membrane Is a Metabolically Active Territory that Generates Nuclear Lipid Droplets. Cell. 174:700–715 e718.

48. Santos-Rosa, H., J. Leung, N. Grimsey, S. Peak-Chew, and S. Siniossoglou. 2005. The yeast lipin Smp2 couples phospholipid biosynthesis to nuclear membrane growth. EMBO J. 24:1931–1941.

49. Soltysik, K., Y. Ohsaki, T. Tatematsu, J. Cheng, and T. Fujimoto. 2019. Nuclear lipid droplets derive from a lipoprotein precursor and regulate phosphatidylcholine synthesis. Nat Commun. 10:473.

50. Tavasoli, M., S. Lahire, T. Reid, M. Brodovsky, and C.R. McMaster. 2020. Genetic diseases of the Kennedy pathways for membrane synthesis. J Biol Chem. 295:17877–17886.

51. Toprakcioglu, Z., and T.P.J. Knowles. 2021. Sequential storage and release of microdroplets. Microsyst Nanoeng. 7:76.

52. van der Veen, J.N., J.P. Kennelly, S. Wan, J.E. Vance, D.E. Vance, and R.L. Jacobs. 2017. The critical role of phosphatidylcholine and phosphatidylethanolamine metabolism in health and disease. Biochim Biophys Acta Biomembr. 1859:1558–1572.

53. van Meer, G., D.R. Voelker, and G.W. Feigenson. 2008. Membrane lipids: where they are and how they behave. Nat Rev Mol Cell Biol. 9:112–124.

54. West, M., N. Zurek, A. Hoenger, and G.K. Voeltz. 2011. A 3D analysis of yeast ER structure reveals how ER domains are organized by membrane curvature. J Cell Biol. 193:333–346.

55. Yang, C., X. Wang, J. Wang, X. Wang, W. Chen, N. Lu, S. Siniossoglou, Z. Yao, and K. Liu. 2020. Rewiring Neuronal Glycerolipid Metabolism Determines the Extent of Axon Regeneration. Neuron. 105:276–292 e275.

56. Zhang, P., L.S. Csaki, E. Ronquillo, L.J. Baufeld, J.Y. Lin, A. Gutierrez, J.R. Dwyer, D.N. Brindley, L.G. Fong, P. Tontonoz, S.G. Young, and K. Reue. 2019. Lipin 2/3 phosphatidic acid phosphatases maintain phospholipid homeostasis to regulate chylomicron synthesis. J Clin Invest. 129:281–295.

